# Human-specific *ARHGAP11B* induces hallmarks of neocortical expansion in developing ferret neocortex

**DOI:** 10.1101/395830

**Authors:** Nereo Kalebic, Carlotta Gilardi, Mareike Albert, Takashi Namba, Katherine R. Long, Milos Kostic, Barbara Langen, Wieland B. Huttner

## Abstract

The evolutionary increase in size and complexity of the primate neocortex is thought to underlie the higher cognitive abilities of humans. *ARHGAP11B* is a human-specific gene that, based on its expression pattern in fetal human neocortex and progenitor effects in embryonic mouse neocortex, has been proposed to have a key function in the evolutionary expansion of the neocortex. Here, we study the effects of *ARHGAP11B* expression in the developing neocortex of the gyrencephalic ferret. In contrast to its effects in mouse, ARHGAP11B markedly increases proliferative basal radial glia, a progenitor cell type thought to be instrumental for neocortical expansion, and results in extension of the neurogenic period and an increase in upper-layer neurons. As a consequence, the postnatal ferret neocortex exhibits an increased neuron density in the upper cortical layers and expands in the radial dimension. Thus, human-specific ARHGAP11B can elicit hallmarks of neocortical expansion in developing ferret neocortex.

## Introduction

The expansion of the neocortex during primate evolution is thought to constitute one important basis for the unparalleled cognitive abilities of humans. The size of the neocortex is mainly regulated by the proliferative potential of neural progenitor cells during cortical development and the length of the neurogenic period (Azevedo et al., 2009; Borrell and Götz, 2014; Dehay et al., 2015; Kaas, 2013; Kalebic et al., 2017; Krubitzer, 2007; Lui et al., 2011; Molnar et al., 2006; Rakic, 2009; Sousa et al., 2017; Wilsch-Bräuninger et al., 2016).

Two major classes of neural progenitors can be distinguished: apical progenitors (APs), whose cell bodies reside in the ventricular zone (VZ), and basal progenitors (BPs), whose cell bodies reside in the subventricular zone (SVZ). Whereas APs are highly proliferative in the neocortex of all mammalian species studied (Götz and Huttner, 2005; Rakic, 2003a), BPs are highly proliferative only in species with an expanded neocortex (Borrell and Götz, 2014; Florio and Huttner, 2014; Lui et al., 2011; Reillo et al., 2011). Specifically, a subtype of BPs, called basal (or outer) radial glia (bRG), are thought to play a key role in the evolutionary expansion of the neocortex (Borrell and Götz, 2014; Florio and Huttner, 2014; Lui et al., 2011). Importantly, in species with an expanded neocortex, such as primates or the ferret, the SVZ has been shown to be divided into two distinct histological zones: the inner and outer SVZ (ISVZ and OSVZ, respectively) (Dehay et al., 2015; Smart et al., 2002). The OSVZ is uniquely important for the evolutionary expansion of the neocortex, as proliferative bRG are particularly abundant in this zone (Betizeau et al., 2013; Fietz et al., 2010; Hansen et al., 2010; Poluch and Juliano, 2015; Reillo et al., 2011; Smart et al., 2002). Increased proliferative capacity of bRG results in an amplification of BP number and is accompanied by a prolonged phase of production of late-born neurons (Geschwind and Rakic, 2013; Otani et al., 2016; Rakic, 2009). As the mammalian cerebral cortex is generated in an inside-out fashion, these late-born neurons occupy the upper-most layers of the cortex (Lodato and Arlotta, 2015; Molnar et al., 2006; Molyneaux et al., 2007; Rakic, 1972, 2009; Sidman and Rakic, 1973). Thus, an increased generation of upper-layer neurons and increased thickness of the upper layers are also hallmarks of an expanded neocortex.

The evolutionary expansion of the neocortex is characteristically accompanied by an increase in the abundance of proliferative bRG, in the length of the neurogenic period, and in the relative proportion of upper-layer neurons within the cortical plate (Borrell and Götz, 2014; Dehay et al., 2015; Florio and Huttner, 2014; Geschwind and Rakic, 2013; Lui et al., 2011; Molnar et al., 2006; Sousa et al., 2017; Wilsch-Bräuninger et al., 2016). This is most obvious when comparing extant rodents, such as mouse, with primates, such as human. Carnivores, such as ferret, display intermediate features (Hutsler et al., 2005; Kawasaki, 2014). Specifically, ferrets exhibit a moderately gyrified neocortex and, during development, a pronounced OSVZ populated with proliferative bRG (Barnette et al., 2009; Borrell and Reillo, 2012; Fietz et al., 2010; Kawasaki, 2014; Kawasaki et al., 2013; Poluch and Juliano, 2015; Reillo et al., 2011; Sawada and Watanabe, 2012; Smart and McSherry, 1986a, b). In this context, it should be noted that in evolution, the split between the lineages leading to mouse and to human occurred a few million years later than that leading to ferret and human (Lewitus et al., 2014). This is consistent with the notion that the features associated with neocortex expansion in extant mammals occurred multiple times in the various lineages of the mammalian phylogenetic tree (Lewitus et al., 2014).

In addition to the above-mentioned features associated with neocortex expansion in general, certain specific aspects of human neocortex expansion are thought to involve human-specific genomic changes. Recent transcriptomic studies established that certain previously identified human-specific genes (Bailey et al., 2002; Dennis and Eichler, 2016) are preferentially expressed in neural progenitor cells and have implicated these genes in human neocortex expansion (Fiddes et al., 2018; Florio et al., 2015; Florio et al., 2018; Florio et al., 2016; Suzuki et al., 2018). Among these genes, the one that showed the most specific expression in human bRG compared to neurons was *ARHGAP11B* (Florio et al., 2015). *ARHGAP11B* arose in evolution after the split of the human lineage from the chimpanzee lineage, as a product of a partial gene duplication of *ARHGAP11A*, a gene encoding a Rho GTPase activating protein (Dennis et al., 2017; Florio et al., 2015; Florio et al., 2016; Kagawa et al., 2013). Forced expression of *ARHGAP11B* in the embryonic mouse neocortex leads to an increase in BP proliferation and pool size (Florio et al., 2015). Here, we study the effects of forced expression of *ARHGAP11B* in the developing ferret neocortex, which already exhibits several features of an expanded neocortex, albeit at an inferior level compared to human.

## Results

We expressed ARHGAP11B in the ferret neocortex starting at embryonic day 33 (E33), when both the generation of upper-layer neurons and formation of the OSVZ start (Martinez-Martinez et al., 2016). Specifically, we performed *in utero* electroporation of ferrets (Kawasaki et al., 2012; Kawasaki et al., 2013) at E33 with a plasmid encoding *ARHGAP11B* under the constitutive CAG promoter or an empty vector as control. The analyses of electroporated embryos were performed at four different developmental stages: E37, E40/postnatal day (P) 0, P10 and P16 (Figure 1 supplement 1A). To be able to visualize the electroporated area, we co-electroporated ARHGAP11B-expressing and control plasmids with vectors encoding fluorescent markers. For postnatal studies, to be able to distinguish the electroporated kits, the ARHGAP11B-expressing plasmid was co-electroporated with a GFP-encoding plasmid, and the control vector with an mCherry-encoding plasmid, or *vice versa*. For the sake of simplicity, we refer to both fluorescent markers as Fluorescent Protein (FP) from here onwards, and both FPs are depicted in green color in all figures.

**Figure 1.**
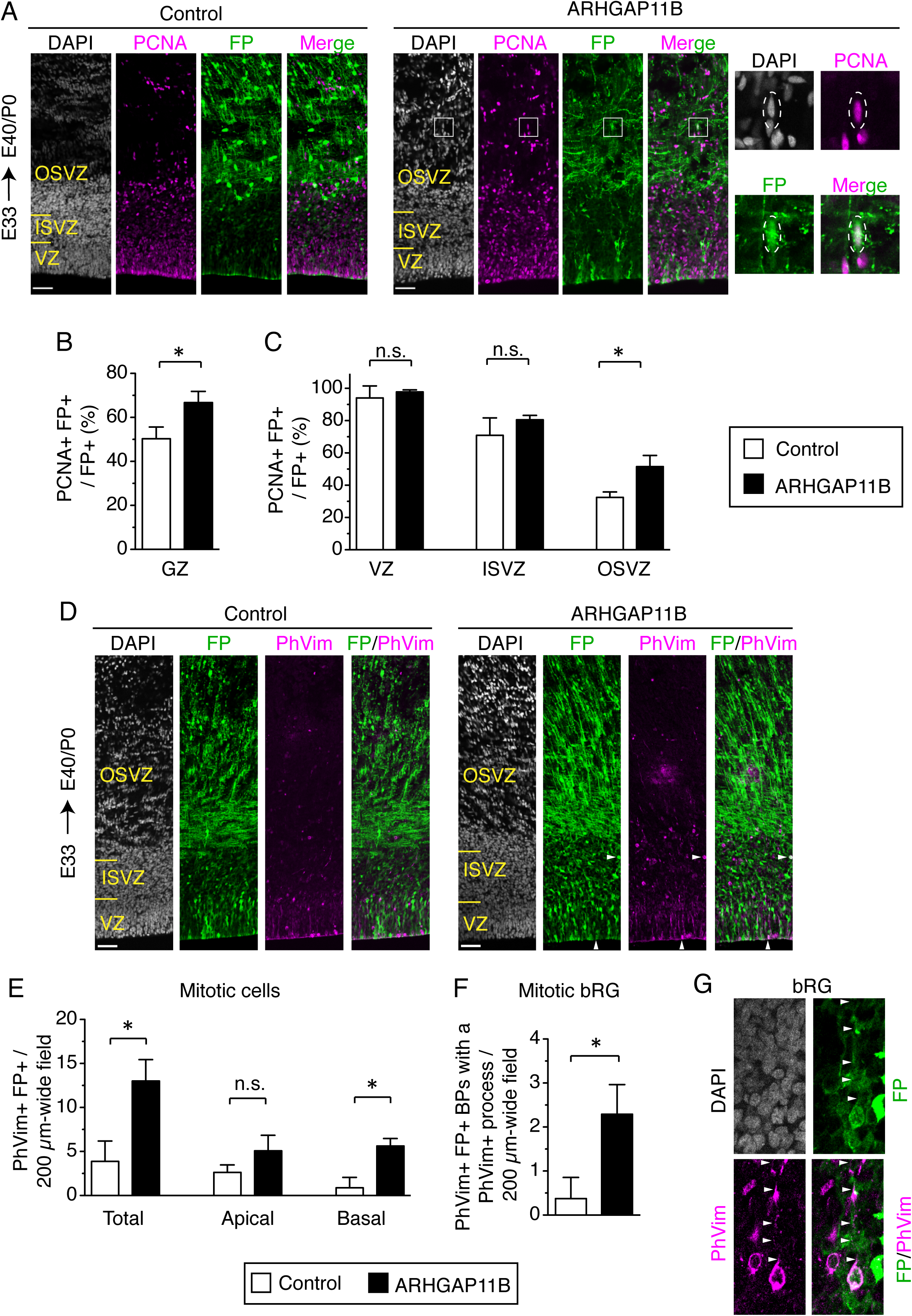
ARHGAP11B increases the abundance of BPs in the developing ferret neocortex. Ferret E33 neocortex was electroporated *in utero* with a plasmid encoding a fluorescent protein (FP) together with either a plasmid encoding ARHGAP11B or empty vector (Control), followed by analysis at E40/P0. (A) Double immunofluorescence for FP (green) and PCNA (magenta), combined with DAPI staining (white). Images are single optical sections. Scale bars, 50 μm. Boxes (50 × 50 μm) indicate an FP+ BP in the OSVZ, shown at higher magnification on the right. Dashed lines indicate a cell body contour. (B, C) Percentage of FP+ cells in the germinal zones (B, GZ) and in the VZ (C, left), ISVZ (C, center) and OSVZ (C, right) that are PCNA+ upon control (white) and ARHGAP11B (black) electroporations. Data are the mean of 3 experiments. Error bars indicate SD; n.s., not statistically significant; *, P <0.05, Student’s *t*-test. (D) Double immunofluorescence for FP (green) and phospho-vimentin (PhVim, magenta), combined with DAPI staining (white). Images are single optical sections. Scale bars, 50 μm. Vertical arrowheads, apical mitosis; horizontal arrowheads, basal mitosis. (E) Quantification of total (left), apical (center) and basal (right) FP+ mitotic cells, as revealed by PhVim immunofluorescence, in a 200 µm-wide field of the cortical wall, upon control (white) and ARHGAP11B (black) electroporations. Data are the mean of 4 experiments. Error bars indicate SD; n.s., not statistically significant, *, P <0.05, Mann-Whitney *U*-test. (F) Quantification of mitotic bRG (FP+ PhVim+ cell bodies in the SVZ that contain a PhVim+ process), in a 200 µm-wide field of cortical wall, upon control (white) and ARHGAP11B (black) electroporations. Data are the mean of 4 experiments. Error bars indicate SD; *, P <0.05, Mann-Whitney *U*-test. (G) Mitotic bRG (single optical sections). Double immunofluorescence for FP (green) and phospho-vimentin (PhVim, magenta), combined with DAPI staining (white), upon electroporation of the plasmid encoding FP together with the plasmid encoding ARHGAP11B. Arrowheads, PhVim+ basal process of the mitotic bRG. Images are oriented with the apical side facing down and are 25 μm wide.

We detected *ARHGAP11B* transcript by RT-qPCR at all the stages analyzed and only in ferret embryos/kits subjected to *ARHGAP11B in utero* electroporation (Figure 1 supplement 1B-D). Additionally, immunofluorescence at E37 demonstrated the specific presence of the ARHGAP11B protein in neural progenitors of such embryos (Figure 1 supplement 1E, F).

## ARHGAP11B increases the abundance of BPs in the developing ferret neocortex

We first examined the ability of ARHGAP11B to increase BP abundance in ferret. To this end, we immunostained E40/P0 ferret neocortex for PCNA, a marker of cycling cells, in order to identify progenitor cells (Figure 1A). We observed an increase in PCNA+ FP+ cells in the OSVZ of the ARHGAP11B-expressing embryos compared to control (Figure 1B, C). We next immunostained the E40/P0 ferret neocortex for phospho-vimentin (PhVim), a marker of mitotic cells (Figure 1D). Our analysis (Figure 1E, left) revealed a 3-fold increase in the abundance of mitotic cells between control and ARHGAP11B-expressing embryos. This increase was 6-fold for mitotic BPs, but only 2-fold (albeit not statistically significant) for mitotic APs (Figure 1E). A comparably large increase as was observed for BPs was detected when examining mitotic bRG, i.e. PhVim+ BPs exhibiting a PhVim+ process (Figure 1F, G), which accounted for 40% of all BPs (compare Figure 1E right, F). Taken together, these data indicate that ARHGAP11B markedly increases the abundance of BP, and specifically of bRG, when expressed in the embryonic ferret neocortex.

## ARHGAP11B increases the proportion of Sox2-positive bRG that are Tbr2-negative

We next analyzed the ARHGAP11B-increased bRG in more detail. Proliferative neural progenitors, in particular apical radial glia (aRG) and bRG, characteristically express the transcription factor Sox2 (Pollen et al., 2015). We therefore immunostained E40/P0 ferret neocortex for Sox2 (Figure 2A) and detected a 40% increase in the proportion of Sox2+ FP+ cells in the germinal zones (GZs) (Figure 2B). This increase was exclusively due to an increase in BPs, as we observed a doubling of the proportion of Sox2+ FP+ cells in both the ISVZ and OSVZ, but no increase in the VZ, upon ARHGAP11B expression (Figure 2C).

**Figure 2.**
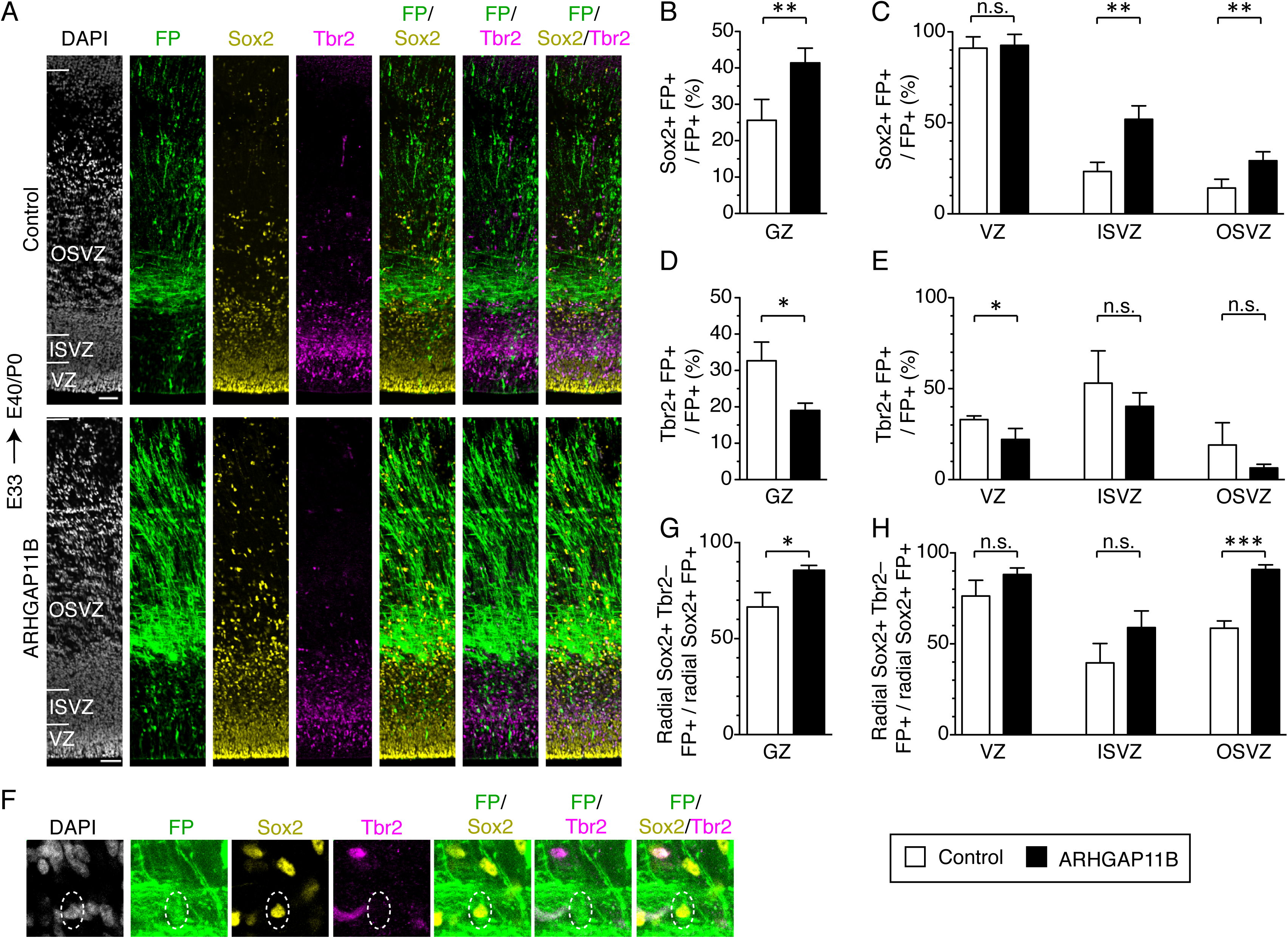
ARHGAP11B increases the proportion of Sox2-positive bRG that are Tbr2-negative. Ferret E33 neocortex was electroporated *in utero* with a plasmid encoding FP together with either a plasmid encoding ARHGAP11B or empty vector (Control), followed by triple immunofluorescence for FP (green), Sox2 (yellow) and Tbr2 (magenta), combined with DAPI staining (white), at E40/P0. (A) Overview of the electroporated areas (single optical sections). Scale bars, 50 μm. (B, C) Percentage of FP+ cells in the germinal zones (B, GZ) and in the VZ (C, left), ISVZ (C, center) and OSVZ (C, right) that are Sox2+ upon control (white) and ARHGAP11B (black) electroporations. (D, E) Percentage of FP+ cells in the germinal zones (D, GZ) and in the VZ (E, left), ISVZ (E, center) and OSVZ (E, right) that are Tbr2+ upon control (white) and ARHGAP11B (black) electroporations. (F) Proliferative bRG (Sox2+ Tbr2– cell in the SVZ exhibiting radial morphology, single optical sections). Triple immunofluorescence for FP (green), Sox2 (yellow) and Tbr2 (magenta), combined with DAPI staining (white), upon electroporation of the plasmid encoding FP together with the plasmid encoding ARHGAP11B. Dashed lines, cell body. Images are oriented with the apical side facing down and are 25 μm wide. (G, H) Percentage of Sox2+ FP+ cells exhibiting radial morphology in the germinal zones (G, GZ) and in the VZ (H, left), ISVZ (H, center) and OSVZ (H, right) that are Tbr2– upon control (white) and ARHGAP11B (black) electroporations. (B-E, G, H) Data are the mean of 4 experiments. Error bars indicate SD; n.s., not statistically significant; ***, P<0.001; **, P <0.01; *, P <0.05, Student’s *t*-test.

We then took advantage of a marker of neurogenic BPs, the transcription factor Tbr2 (Figure 2A) (Englund et al., 2005), which is expressed in mouse bRG, but not proliferative human bRG (Fietz et al., 2010; Florio et al., 2015; Hansen et al., 2010; Pollen et al., 2015). Upon expression of ARHGAP11B for 7 days, we observed a decrease in the proportion of Tbr2+ FP+ progenitors in all GZs, which was statistically significant for the VZ (Figure 2D, E).

In order to potentially obtain further cues as to the proliferative potential of the ARHGAP11B-increased bRG in the developing ferret neocortex, we focused our attention on progenitor cells that (i) exhibited a radial morphology, (ii) expressed Sox2, but (iii) lacked Tbr2 expression (Figure 2F). Upon ARHGAP11B expression, we observed, in the sum of the GZs, a 20% increase in the proportion of radial Sox2+ cells that were Tbr2– (Figure 2G). This was largely due to an increase in the proportion of these cells in the OSVZ (Figure 2H), where more than 90% were Tbr2–. These findings are consistent with the notion that the ARHGAP11B-increased radial Sox2+ Tbr2– cells in the OSVZ are bRG. Studies in fetal human neocortex have established that such cells in the OSVZ are highly proliferative ((Hansen et al., 2010; LaMonica et al., 2013) for reviews see (Florio and Huttner, 2014; Lui et al., 2011)). Hence, our finding that expression of ARHGAP11B in developing ferret neocortex results in a marked increase in the proportion of these cells suggests that this human-specific gene is sufficient to promote, in a gyrencephalic carnivore, the generation of bRG with increased proliferative potential.

## ARHGAP11B expression in developing ferret neocortex results in an extended neurogenic period

We investigated the potential consequences of the ARHGAP11B-elicited increase in the abundance of BPs, notably of Sox2+ Tbr2– bRG, for neurogenesis in the developing ferret neocortex. To this end, we immunostained E40/P0 ferret neocortex for Tbr1, a transcription factor which is a marker of deep-layer neurons (Kolk et al., 2006) (Figure 3 supplement 1A), and for Satb2, a transcriptional regulator which is expressed in neurons that establish callosal projections and that are highly enriched in the upper layers of the cortical plate (CP) (Alcamo et al., 2008; Britanova et al., 2008) (Figure 3A). The vast majority (>90%) of the FP+ neurons in the CP of both control and ARHGAP11B-expressing ferret neocortex were found to be Satb2+ (Figure 3 supplement 1B). This high percentage is consistent with our experimental approach in which we targeted the embryonic ferret neural progenitors by *in utero* electroporation at E33, i.e. the time when the generation of the upper-layer neurons starts (Jackson et al., 1989; Martinez-Martinez et al., 2016).

**Figure 3.**
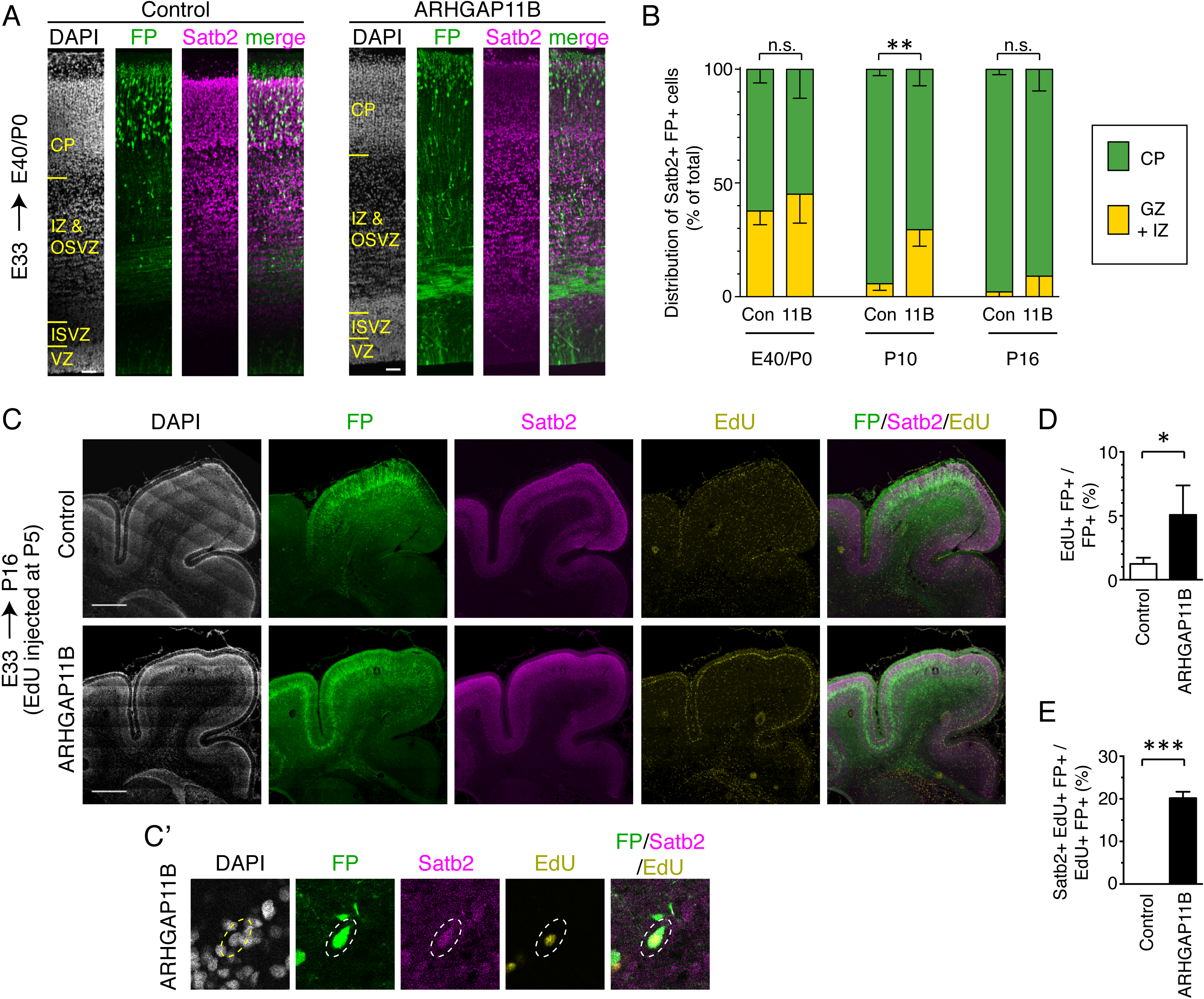
ARHGAP11B expression in developing ferret neocortex results in an extended neurogenic period. Ferret E33 neocortex was electroporated *in utero* with a plasmid encoding FP, together with either a plasmid encoding ARHGAP11B or empty vector (Control), followed by analysis at E40/P0 (A, B left), P10 (B center) and P16 (B right, C-E). (A) Double immunofluorescence for FP (green) and Satb2 (magenta), combined with DAPI staining (white), of the E40/P0 ferret neocortex. The immunofluorescence of the same cryosection for Tbr1 is shown in Figure 3 supplement 1A. Images are single optical sections. Scale bars, 50 μm. (B) Distribution of Satb2+ FP+ neurons at E40/P0 (left), P10 (center) and P16 (right), between the cortical plate (CP, green) and germinal zones plus intermediate zone (GZ+IZ, yellow), upon control (Con, left) and ARHGAP11B (11B, right) electroporations. Data are the mean of 3 (P0 and P10) or 4 (P16) experiments. Error bars indicate SD; **, P <0.01; n.s., not statistically significant, two-way ANOVA with Bonferroni post-hoc tests (P10, Control CP vs. ARHGAP11B CP, P=0.0015). (C) Triple (immuno)fluorescence for FP (green), Satb2 (magenta) and EdU (yellow), combined with DAPI staining (white), of the P16 ferret neocortex, upon EdU injection at P5. Images are single optical sections. Scale bars, 1 mm. (C’) Higher magnification of a FP+ Satb2+ EdU+ neuron upon electroporation of the plasmid encoding FP together with the plasmid encoding ARHGAP11B. Dashed lines, cell body. Images (single optical sections) are oriented with the apical side facing down and are 50 μm wide. (D) Percentage of FP+ cells that are EdU+ upon control (white) and ARHGAP11B (black) electroporations. (E) Percentage of EdU+ FP+ cells that are Satb2+ upon control (white) and ARHGAP11B (black) electroporations. (D, E) Data are the mean of 3 experiments. Error bars indicate SD; ***, P <0.001; *, P <0.05, Student’s *t*-test.

Analysis of the distribution of Satb2+ FP+ neurons at E40/P0 between the CP on the one hand side, and the GZs plus the intermediate zone (IZ) on the other hand side, revealed that around 60% of the neurons had reached the CP in both control and ARHGAP11B-expressing ferret neocortex (Figure 3B left). We next examined electroporated ferret neocortex at P10 (Figure 3 supplement 2; analysis confined to gyri), which is the stage when neuron production is completed and neuron migration is terminating in the motor and somatosensory areas (Jackson et al., 1989; Smart and McSherry, 1986a, b). Consistent with this, our analysis of the control brains revealed that more than 90% of the Satb2+ FP+ neurons had reached the CP (Figure 3B middle). In contrast, only 70% of the Satb2+ FP+ neurons were found in the CP of the ARHGAP11B-expressing neocortex (Figure 3B middle). However, at P16, nearly all Satb2+ FP+ neurons were found in the CP in both control and ARHGAP11B-expressing neocortex (Figure 3B right; again, analysis confined to gyri). These data suggested that upon ARHGAP11B expression, either neurons migrate more slowly to the CP, or the neurogenic period is extended.

To explore the latter scenario, we injected EdU into P5 ferret kits (i.e. 12 days after electroporation), which is the stage when neuronal progenitors undergo their very last neuron-generating cell divisions in the motor and somatosensory areas of the neocortex (Jackson et al., 1989; Smart and McSherry, 1986a, b). Analysis 11 days after EdU injection, at P16 (Figure 3C), revealed a 4-fold increase in the proportion of FP+ cells that were EdU+, in neocortical gyri of ARHGAP11B-expressing kits compared to control (Figure 3D). This was consistent with a prolonged, and hence increased, production of cells in ARHGAP11B-expressing kits, which in turn would be in line with the above described finding that ARHGAP11B increases the abundance of proliferative bRG. Importantly, among the EdU+ FP+ cells of the ARHGAP11B-expressing neocortex, 20% were Satb2+ neurons (Figure 3C’ and E). In contrast, we did not detect a single Satb2+ EdU+ FP+ neuron in any of the control neocortices (Figure 3E). Collectively, these data indicate that neurogenesis in ARHGAP11B-expressing ferret neocortex continues longer than in control neocortex.

## ARHGAP11B expression in developing ferret neocortex results in a greater abundance of upper-layer neurons

In light of the extension of the neurogenic period upon ARHGAP11B expression, we examined a potential increase in the abundance of the last-born neurons, that is, the upper-layer neurons. To this end, we first performed Nissl staining of the P16 ferret neocortex to visualize all neurons and the various layers of the CP, and immunostaining for Satb2, which is expressed in the majority of upper-layer neurons (Alcamo et al., 2008; Britanova et al., 2008; Lodato and Arlotta, 2015) (Figure 4A). These analyses, carried out on gyri, revealed (i) an increase in the abundance of FP+ cells in the CP (Figure 4 supplement 1A), and (ii) an alteration in the distribution of the FP+ cells between layers II-VI of the CP, with a greater proportion of cells in layers II-IV (Figure 4 supplement 1B), in the ARHGAP11B-expressing neocortex compared to control. A similar abundance increase (Figure 4 supplement 1C) and altered distribution (Figure 4B) were observed for Satb2+ FP+ neurons. Furthermore, the proportion of FP+ cells in the CP that were Satb2+ neurons was increased upon ARHGAP11B expression (Figure 4C).

**Figure 4.**
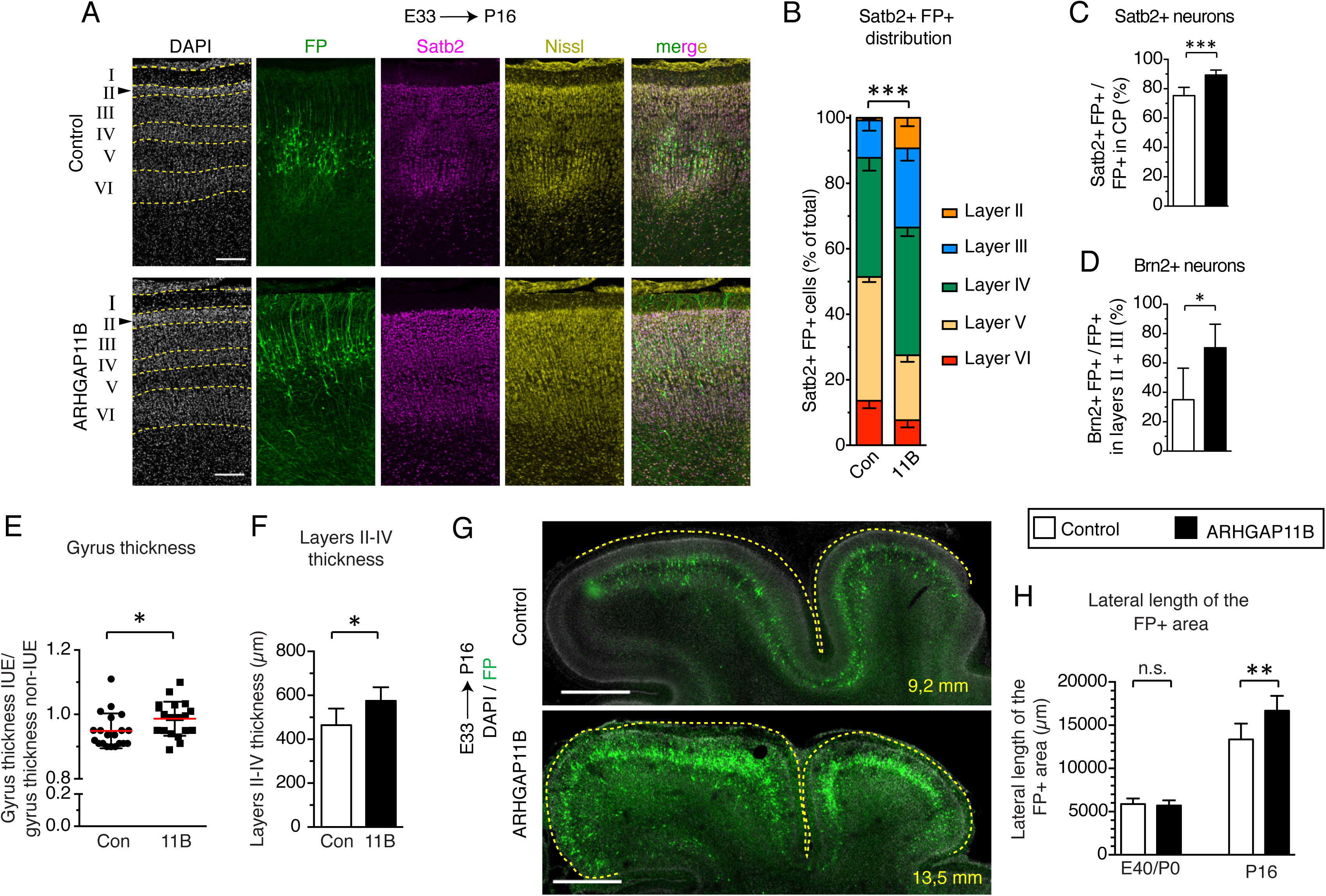
ARHGAP11B expression results in a greater abundance of upper-layer neurons and expansion of the developing ferret neocortex. Ferret E33 neocortex was electroporated *in utero* with a plasmid encoding FP, together with either a plasmid encoding ARHGAP11B or empty vector (Control), followed by analysis at E40/P0 (G, left) or P16 (all other panels). (A) Double immunofluorescence for FP (green) and Satb2 (magenta), combined with DAPI (white) and Nissl (yellow) staining, of the CP (single optical sections). Neuronal layers are marked on the left. Arrowheads, increased thickness of layer II upon ARHGAP11B expression. Scale bars, 200 μm. (B) Distribution of Satb2+ FP+ neurons between the neuronal layers upon control (Con, left) and ARHGAP11B (11B, right) electroporations. Data are the mean of 6 experiments. Error bars indicate SD; ***, P <0.001, two-way ANOVA with Bonferroni post-hoc tests (Layer V, Control vs. ARHGAP11B, P <0.0001; Layer III, Control vs. ARHGAP11B, P=0.0073) (C) Percentage of FP+ cells in the CP that are Satb2+, upon control (white) and ARHGAP11B (black) electroporations. Data are the mean of 6 experiments. Error bars indicate SD; ***, P <0.001, Student’s *t*-test. (D) Percentage of FP+ cells in layers II+III that are Brn2+, upon control (white) and ARHGAP11B (black) electroporations. Data are the mean of 5 experiments. Error bars indicate SD; *, P <0.05, Student’s *t*-test. (E)Quantification of the gyrus thickness of control (Con) and ARHGAP11B-expressing (11B) ferret neocortex. Measurements were performed as described in Figure 4 supplement 3. All data are expressed as ratio between electroporated hemisphere (IUE) and non-electroporated contralateral hemisphere (non-IUE). Data are the mean (red lines) of 20 gyri per condition from 6 neocortices per condition. Error bars indicate SD; *, P <0.05, Student’s *t*-test. (F) Quantification of layers II-IV thickness, upon control (Con, white) and ARHGAP11B (11B, black) electroporations. Data are the mean of 6 experiments. Error bars indicate SD; *, P <0.05, Student’s *t*-test. (G) Immunofluorescence for FP (green), combined with DAPI staining (white), of the areas harbouring FP+ cells (single optical sections). Dashed lines, basal contour of the FP+ area. Lateral length of the respective contour is indicated in the bottom right corner. Note that the images and the countours depict only the areas with a high abundance of FP+ cells, which is distinct from the quantification shown in (G). Scale bars, 1 mm. (H) Quantification of the lateral length of the entire areas harbouring FP+ cells, measured as depicted in Figure 4 supplement 3D middle, at E40/P0 (left) and P16 (right), upon control (Con, white) and ARHGAP11B (11B, black) electroporations. Data are the mean of 3 (E40/P0) and 6 (P16) experiments. Error bars indicate SD; **, P <0.01, Student’s *t*-test.

We sought to corroborate these data by immunostaining the P16 ferret neocortex for Brn2 (Pou3f2, Figure 4 supplement 2A), a transcription factor considered to be a marker of layer II and III neurons (Dominguez et al., 2013; McEvilly et al., 2002; Sugitani et al., 2002). Quantification of Brn2+ FP+ cells in layers II and III of gyri revealed a marked increase in the abundance of these cells (Figure 4 supplement 2B) and in the proportion of FP+ cells that were Brn2+ neurons (Figure 4D), in ARHGAP11B-expressing kits compared to control. Of note, the effects of ARHGAP11B expression on the Brn2+ neurons were of a greater degree than those on the Satb2+ neurons (compare Figure 4 supplement 2B with Figure 4 supplement 1C; Figure 4, panel D with panel C), in line with the notion that Brn2 exhibits a greater specificity for layer II and III neurons than Satb2 (Dominguez et al., 2013; Sugitani et al., 2002). Taken together, our data indicate that the extension of the neurogenic period upon ARHGAP11B expression in the developing ferret neocortex is accompanied by an increase in the abundance of layer II and III neurons.

## ARHGAP11B expression in developing ferret neocortex leads to its radial expansion

Finally, we explored whether the ARHGAP11B-elicited increase in the abundance of BPs, notably of proliferative bRG, resulting in an extended neurogenic period and in an increased abundance of upper-layer neurons, had any consequences for the size and morphology of the ferret neocortex. To this end, we analyzed the neocortex at P16 (Figure 4 supplement 3A-C), the developmental stage when cortical folds have already formed in the ferret (Barnette et al., 2009; Fernandez et al., 2016; Matsumoto et al., 2017; Sawada and Watanabe, 2012; Shinmyo et al., 2017), and quantified a set of morphological parameters (Figure 4 supplement 3D), each on two coronal sections located slightly rostrally (Figure 4 supplement 3B and C, position 1) and slightly caudally (Figure 4 supplement 3B and D, position 2), respectively, to a middle position along the rostro-caudal axis. This revealed no significant differences between control and ARHGAP11B-expressing kits with regard to brain mass (Figure 4 supplement 3E) and gross neocortex morphology, including the gyrification index of the electroporated area (referred to as local GI), the size of the posterior sigmoid gyrus, lateral gyrus and coronal gyrus, and the depth and thickness of the cruciate sulcus, suprasylvian sulcus and lateral sulcus (Figure 4 supplement 3F-I).

However, we did detect an increase, upon ARHGAP11B expression, in the thickness of the three gyri analyzed (Figure 4E). We therefore investigated the underlying cause of this thickness increase and, considering the increased abundance of upper-layer neurons upon ARHGAP11B expression (Figure 4 supplement 1C, Figure 4 supplement 2B), measured the thickness of CP layers II-IV in the gyri of control and ARHGAP11B-expressing ferret neocortex at P16. This revealed a 100-µm increase in the thickness of these upper layers in ARHGAP11B-expressing neocortex compared to control (Figure 4F, note also the thicker layer II in Figure 4A and Figure 4 supplement 3K). Hence, ARHGAP11B is sufficient to promote neocortex expansion in ferret in the radial dimension.

We then explored whether ARHGAP11B would also be able to promote neocortex expansion in ferret in the tangential dimension. Here, we exploited the previous findings that during the first week of the ferret postnatal neocortex development, migrating neurons change their migration mode from radial to tangential, which results in an increased lateral dispersion of the late-born neurons (Gertz and Kriegstein, 2015; Reillo et al., 2011; Smart and McSherry, 1986a, b). In line with this, the abundance of FP+ cells per 200 µm-wide field of cortical wall decreased about 3-fold from E40/P0 to P16 (Figure 4 supplement 3J). Yet, the increase in FP+ cells per field of cortical wall due to ARHGAP11B expression was not only observed at P16, confirming the above described analysis of FP+ cell numbers in the CP (Figure 4 supplement 1A), but was already detected at E40/P0 (Figure 4 supplement 3J). The decrease in FP+ cell abundance per 200 µm-wide field of cortical wall from E40/P0 to P16 (Figure 4 supplement 3J) was not accompanied by an increase in cell death (Figure 4 supplement 4). Rather, this decrease reflected an increase in the lateral spread of the progeny of the targeted cells, as evidenced by the ARHGAP11B-elicited increase in the lateral length of the area harbouring FP+ cells that was observed at P16, but not at E40/P0 (Figure 4G,H). Hence, ARHGAP11B is sufficient to affect neocortex development in ferret also in the tangential dimension.

However, given that ARHGAP11B expression in the developing ferret neocortex increased the lateral length of the area harbouring FP+ cells but did not increase gyrus size, we explored whether this discrepancy could be resolved by ARHGAP11B expression increasing cell density, specifically in the upper layers of the CP where most of the ARHGAP11B-increased neurons resided. Indeed, our analysis revealed an increase in cell density in layers III and IV at P16, but not in layer II, which in general exhibited the highest cell density (Figure 4 supplement 3K, L). Taken together, these data indicate that the ARHGAP11B-promoted increase in upper-layer neurons, which resulted in their greater lateral spread, led to a higher cell density in those upper layers that were able to accommodate additional neurons, rather than to an increased gyrus size.

## ARHGAP11B induces the appearance of astrocytes from the targeted progenitors in the developing ferret neocortex

Neocortex expansion involves not only an increase in the number of neurons, but also of glial cells. Although we on purpose performed *in utero* electroporation of the ferret neocortex at E33 in order to target OSVZ progenitors generating upper-layer neurons (Jackson et al., 1989; Martinez-Martinez et al., 2016), and therefore would expect only a small proportion of the FP+ cells to be glial cells, we nonetheless examined the possible occurrence of GFAP+ and S100ß+ cells (as indicators of an astrocytic lineage (Reillo et al., 2011; Voigt, 1989)) and of Olig2+ cells (as an indicator of an oligodendrocytic lineage (Mo and Zecevic, 2009; Voigt, 1989)) among the FP+ progeny of the targeted cells. An additional reason to examine astrocytes was that radial glial progenitor cells at later stages of human, monkey and ferret neocortex development are known to differentiate into astrocytic progenitors which then give rise to post-mitotic astrocytic cells (Rakic, 1978, 2003b; Reillo et al., 2011; Voigt, 1989).

Upon immunostaining of the P16 ferret neocortex for GFAP (Figure 4 supplement 5A, B) and Olig2 (Figure 4 supplement 5E), no FP+ cells in the CP of control neocortex that were GFAP+ or Olig2+ could be detected. In contrast, while we still did not detect any Olig2+ FP+ cells in the CP of ARHGAP11B-expressing neocortex, about 3% of the CP FP+ cells were GFAP+ (Figure 4 supplement 5C). Given that ARHGAP11B expression increases the pool size of FP+ cells (Figure 4 supplement 1A, Figure 4 supplement 3J), this translates into an ARHGAP11B-induced appearance of GFAP+ FP+ cells in the CP. All of these GFAP+ FP+ cells were Ki67– (data not shown) and hence post-mitotic astrocytes. To corroborate these data, we performed immunostaining of the P16 ferret neocortex for another astrocyte marker, S100ß (Lopez-Hidalgo et al., 2016), and readily identified FP+ S100ß+ astrocytes upon ARHGAP11B expression. These data are consistent with at least two scenarios related to the increased abundance of proliferative bRG upon ARHGAP11B expression. Either this increase results not only in an extension of the neurogenic period but also in the initiation of astrogenesis. Or the greater bRG pool size gives rise to a detectable level of astrocytic cells once the bRG differentiate along the astrocytic lineage (Rakic, 2003b; Reillo et al., 2011; Voigt, 1989).

## Discussion

We have shown that expression of a single human-specific gene, *ARHGAP11B*, which has been implicated in the evolutionary expansion of the human neocortex (Florio et al., 2015; Florio et al., 2016), is sufficient to elicit features in the developing ferret neocortex that are characteristically associated with a further expanded neocortex such as human. These features include (i) an increase in the pool size of proliferative BPs, notably proliferative bRG; (ii) a lengthening of the neurogenic period; (iii) an increased generation of neurons, in particular of upper-layer neurons; and (iv) an increase in the size of the CP that harbours these additional neurons, reflected by the greater thickness of its upper layers and the larger lateral area where these neurons are found.

### Increase in Proliferative bRG

The present effects of ARHGAP11B on BPs in developing ferret neocortex are substantially greater than, and significantly different from, those previously observed upon transient *ARHGAP11B* expression in embryonic mouse neocortex (Florio et al., 2015), with regard to BP quantity and quality. As to quantity, in embryonic mouse neocortex, *ARHGAP11B* expression resulted in a doubling and tripling of the pool size of total BPs and BPs in mitosis, respectively (Florio et al., 2015), whereas in embryonic ferret neocortex, *ARHGAP11B* expression led to a 3.5-fold and 6-fold increase in the pool size of total BPs and BPs in mitosis (notably mitotic bRG), respectively (for a data summary, see Supplemental Table 1). The greater effect of *ARHGAP11B* expression on mitotic BP abundance than total BP abundance, in both embryonic mouse and ferret neocortex, indicates that *ARHGAP11B* expression increases the proportion of the total duration of the BP cell cycle that is used for M-phase. This in turn would be consistent with ARHGAP11B promoting, in embryonic ferret neocortex, the proliferative rather than neurogenic mode of bRG division. This notion would be in line with the previous findings that both, cortical progenitors in embryonic mouse neocortex (Arai et al., 2011) and radial glial progenitors in postnatal ferret neocortex (Turrero Garcia et al., 2015), which are not yet committed to neurogenesis spend a greater proportion of their total cell cycle in M-phase than those committed to neurogenesis.

As to BP quality, mouse bRG have been shown to undergo mostly neurogenic cell divisions and to display a limited proliferative capacity (Shitamukai et al., 2011; Wang et al., 2011; Wong et al., 2015). In contrast, human bRG are known to be highly proliferative, which is thought to be one of the key factors contributing to the evolutionary expansion of the human neocortex (Fietz et al., 2010; Florio and Huttner, 2014; Hansen et al., 2010; Lui et al., 2011). The proliferative capacity of bRG in other mammalian species that have been studied appears to follow the general rule that highly proliferative bRG are more often found in species with an expanded neocortex (Betizeau et al., 2013; Reillo et al., 2011). Hence, the ARHGAP11B-promoted increase in the number of proliferative (Sox2+ Tbr2–) bRG in the developing ferret neocortex fulfills a key requirement for further neocortex expansion.

In this context, it should be emphasized that those of the human-specific genes (Dennis and Eichler, 2016) that have been shown to be preferentially expressed in cortical progenitor cells (Florio et al., 2018) and that have been ectopically expressed in embryonic mouse neocortex, i.e. *ARHGAP11B* (Florio et al., 2015) and *NOTCH2NL* (Fiddes et al., 2018; Florio et al., 2018; Suzuki et al., 2018), show an increase in BP proliferation that mostly involves basal intermediate progenitors. To the best of our knowledge, our study provides first evidence of a human-specific gene increasing the number of proliferative bRG in a gyrencephalic neocortex. As bRG are considered to be instrumental for the evolutionary expansion of neocortex, this suggests that the ferret likely is a model system superior to mouse for studying the role of human-specific genes in neocortex development.

### Increase in Upper-Layer Neurons

An increase in the upper layers is a fundamental characteristic of neocortex expansion (Fame et al., 2011; Hutsler et al., 2005; Molnar et al., 2006; Tarabykin et al., 2001). Layers II-IV are evolutionarily the most novel and mammalian-specific layers (Molnar et al., 2006). Layers II and III, in particular, underwent disproportional expansion during primate evolution. As the layers II-III neurons are the last-born neurons, an increase in the length of the neurogenic period might result in an increase in the number of late-born neurons. In fact, it has been proposed that the length of the neurogenic period may be a contributing factor in explaining the evolutionary increase in neocortex size and neuron number between humans and other great apes (Lewitus et al., 2014). Consistent with this, we observed a prolonged generation of late-born neurons, increased thickness of layers II-IV, and a marked increase in Satb2+ (3.3-fold) and, in particular, Brn2+ (5.6-fold) neurons in the upper layers of the CP, upon *ARHGAP11B* expression (for a data summary, see Supplemental Table 1). Thus, also with regard to the key product generated by cortical progenitor cells, i.e. the neurons, ARHGAP11B is sufficient to promote another feature in the developing ferret neocortex that would be consistent with further neocortex expansion.

In this context, Satb2+ neurons are of special interest, as Satb2 is essential for establishing callosal projections to the contralateral hemisphere (Alcamo et al., 2008; Britanova et al., 2008). Callosal projection neurons are considered to play a key role in the high-level associative connectivity, thus contributing significantly to human cognitive abilities, with their impairment causing cognition-related pathologies (Fame et al., 2011). Given that the evolutionary increase in neocortex size is accompanied by an increase in callosal projection neurons, our data show that ARHGAP11B elicits yet another, specific feature in the developing ferret neocortex associated with further neocortex expansion.

### Neocortex expansion vs. increased CP Thickness and Lateral Area

A hallmark of neocortex expansion is neocortical folding (Borrell, 2018; Borrell and Götz, 2014; Kroenke and Bayly, 2018). Transient expression of *ARHGAP11B* in the embryonic mouse neocortex induced cortical folding in about half of the cases in this normally lissencephalic rodent (Florio et al., 2015). However, expression of ARHGAP11B in the developing ferret neocortex did not significantly increase the GI, gyrus size, or sulcus depth and thickness in this moderately gyrencephalic carnivore. The major morphological signs of neocortical expansion upon *ARHGAP11B* expression in developing ferret neocortex that we could detect were (i) an increased gyrus thickness, which reflected the already discussed increase in upper-layer thickness due to the increase in upper-layer neurons, and (ii) a larger lateral area of the CP where these neurons were found. The latter phenotype, however, did not result in an increase in gyrus size.

The reason why the increased upper-layer neurons generated upon *ARHGAP11B* expression – although increasing their final destination CP area in the tangential dimension – did not cause an increase in gyrus width, was that these neurons added themselves to the neurons already existing in the control condition, thereby increasing neuronal cell density in the upper layers. Analysis of cortical neuron density as a function of cortical neuron number across carnivores has revealed that neuron density tends to decrease with increasing neuron number (Herculano-Houzel et al., 2015; Jardim-Messeder et al., 2017; Lewitus et al., 2012). This trend is much less prominent in primates, which in general exhibit a greater cortical neuron density per cortical neuron number than carnivores. Thus, the increase in upper-layer neuron density upon increasing upper-layer neuron number by the human-specific *ARHGAP11B* can be regarded as a form of “primatization” of the ferret neocortex.

In this context, it is interesting to note that neocortical neuron density across carnivores shows a much greater range of variation than that across primates. We therefore hypothesize that the ferret neocortex is plastic enough to accommodate additional neurons, without having to increase neocortex size or folding. A corollary of this notion with regard to ARHGAP11B is that its primary function is to increase the proliferation and thus pool size of BPs, notably of bRG, which results in a lengthening of the neurogenic period and an increased generation of neurons, in particular of upper-layer neurons. Whether this leads to an increase in cortical folding as an indicator of neocortex expansion, as found in ARHGAP11B-expressing mouse embryos (Florio et al., 2015), or to an increase in cortical neuron density, as observed in the present study with ARHGAP11B-expressing ferret kits, appears to reflect more the inherent properties of the mammalian order under study how to deal with an increase in cortical neuron number, rather than the primary function of ARHGAP11B.

## Author contributions

Conceptualization, N.K., W.B.H.; Formal Analysis, N.K.; Investigation, N.K., C.G., M.A.; Methodology, N.K., C.G., K.R.L., M.K., B.L.; Visualization, N.K., C.G.; Resources, T.N.; Writing - Original draft, N.K.; Writing - Review & editing, N.K., W.B.H.; Supervision, W.B.H.; Project Administration and Funding Acquisition, W.B.H.. All authors gave input on the manuscript.

## Acknowledgements

We are grateful to the Services and Facilities of the Max Planck Institute of Molecular Cell Biology and Genetics for the outstanding support provided, notably J. Helppi and his team of the Biomedical services (BMS) for the excellent husbandry of ferrets, J. Peychl and his team of the Light Microscopy Facility and P.Keller and his team of the Antibody Facility. We would like to particularly thank Anke Münch-Wuttke for help with perfusion of ferret kits and Dr. Anna Pfeffer for exceptional veterinary support. NK was supported by an EMBO long-term fellowship (ALTF 861-2013). CG acknowledges support from the Erasmus+ traineeship program and MA from the Christiane-Nüsslein-Volhard Foundation. WBH was supported by grants from the DFG (SFB 655, A2), the ERC (250197) and ERA-NET NEURON (MicroKin).

## Supplemental figure legends

**Figure 1 supplement 1.**
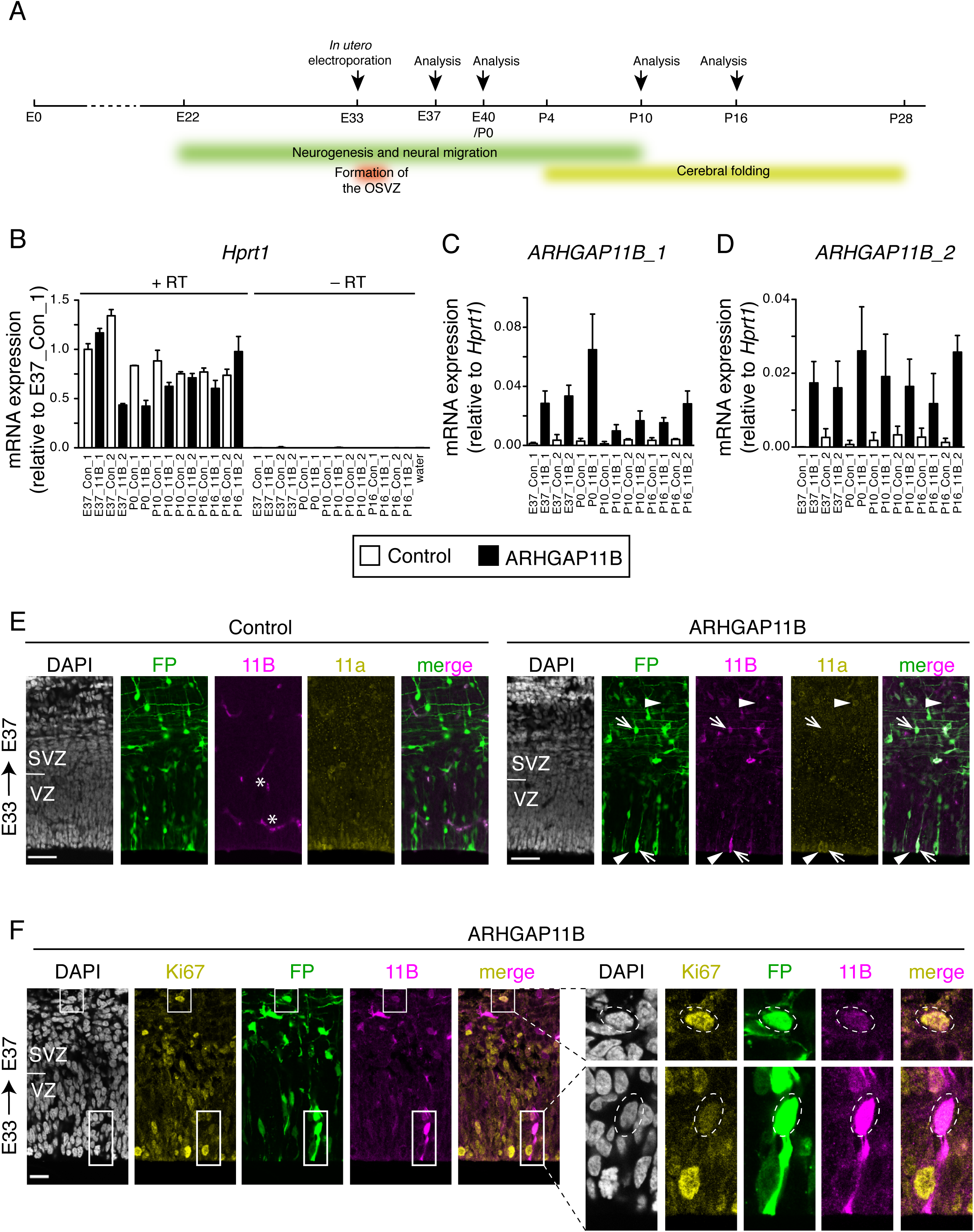
Forced expression of ARHGAP11B in the developing ferret neocortex. (A) Scheme of ferret neocortex development and experimental approach. ARHGAP11B was expressed in developing ferret neocortex by *in utero* electroporation at E33, when the OSVZ begins to form. Analyses were performed at E37, E40/P0, P10 and P16. Analysis of cortical progenitors was performed at E40/P0, analysis of post-mitotic cells at E40/P0, P10 and P16, and analysis of brain size and neocortex morphology at P16. (B-D) mRNA expression analysis by RT-qPCR at E37, P0, P10 and P16. RNA was isolated from cryosections of paraformaldehyde-fixed brain tissue following ferret *in utero* electroporation at E33. Expression of the housekeeping gene *Hprt1* (B, note the lack of signal in the absence of reverse transcriptase (–RT)) was used for normalization of *ARHGAP11B* expression detected with two different primer pairs (C, *ARHGAP11B_1*; D, *ARHGAP11B_2*). Two control (Con_1, Con_2; white) and two ARHGAP11B-electroporated (11B_1, 11B_2; black) embryos were analyzed at each stage, except for P0 when one control and one ARHGAP11B-electroporated embryo were analyzed. Error bars represent SD of three PCR amplifications. (E, F) ARHGAP11B protein expression analysis by immunofluorescence. Ferret E33 neocortex was electroporated *in utero* with a plasmid encoding FP together with either a plasmid encoding ARHGAP11B or empty vector (Control), followed by analysis at E37. (E) Triple immunofluorescence for FP (green), ARHGAB11B (11B, magenta) and Arhgap11a (11a, yellow), combined with DAPI staining (white). Arrows, an ARHGAP11B+ FP+ cell that is Arhgap11a–; arrowheads, an Arhgap11a+ cell that is FP– ARHGAP11B–. Note that the exposure of images of ARHGAP11B staining in the control neocortex was longer than in the ARHGAP11B-electroporated neocortex, in order to show the lack of a specific signal in the control, where only unspecific signal at blood vessels was detected (asterisks). Images are single optical sections. Scale bars, 50 μm. (F) Triple immunofluorescence for FP (green), ARHGAB11B (11B, magenta) and Ki67 (yellow), combined with DAPI staining (white), upon electroporation of a plasmid encoding FP and a plasmid encoding ARHGAP11B. Images are single optical sections. Scale bar, 20 μm. Boxes (25 μm wide), indicating an ARHGAP11B-expressing BP (upper box) and an ARHGAP11B-expressing AP (lower box), are shown at higher magnification on the right. Dashed lines, cell bodies.

**Figure 3 supplement 1.**
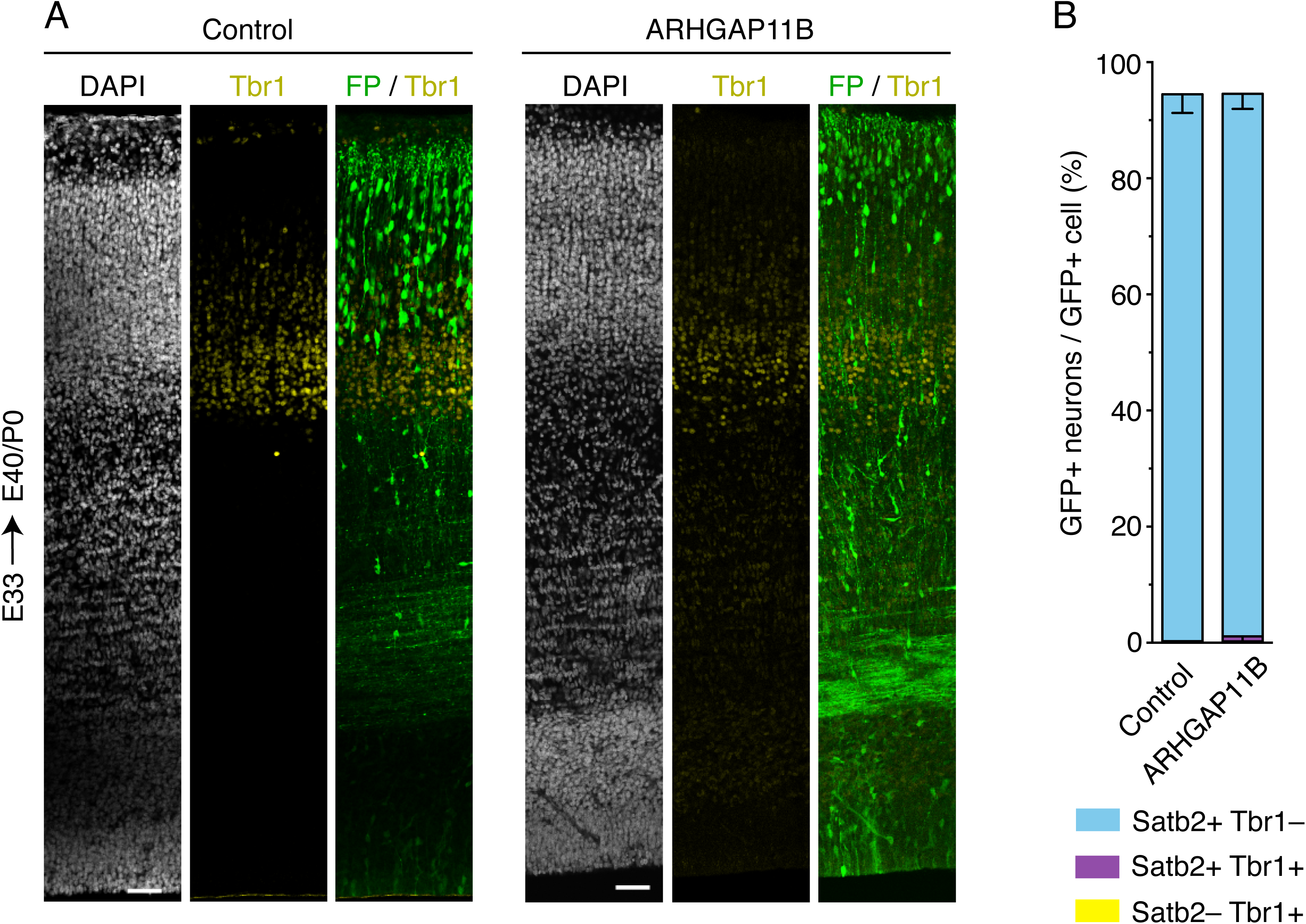
Almost all neurons generated from ARHGAP11B-expressing progenitors are Satb2+. Ferret E33 neocortex was electroporated *in utero* with a plasmid encoding FP together with either a plasmid encoding ARHGAP11B or empty vector (Control), followed by triple immunofluorescence for FP, Tbr1 and Satb2, combined with DAPI staining, at E40/P0. (A) Overview of the electroporated areas showing the immunofluorescence for Tbr1 (yellow) and FP (green) and the DAPI staining (white) (single optical sections). Note that the DAPI staining and FP immunofluorescence images are the same as in Figure 3A; for Satb2 staining, see Figure 3A. Scale bars, 50 μm. (B) Percentages of FP+ cells that are Satb2+ Tbr1– (blue), Satb2+ Tbr1+ (purple) and Satb2– Tbr1+ (yellow), upon control (left) and ARHGAP11B (right) electroporations. Data are the mean of 3 experiments. Error bars indicate SD.

**Figure 3 supplement 2.**
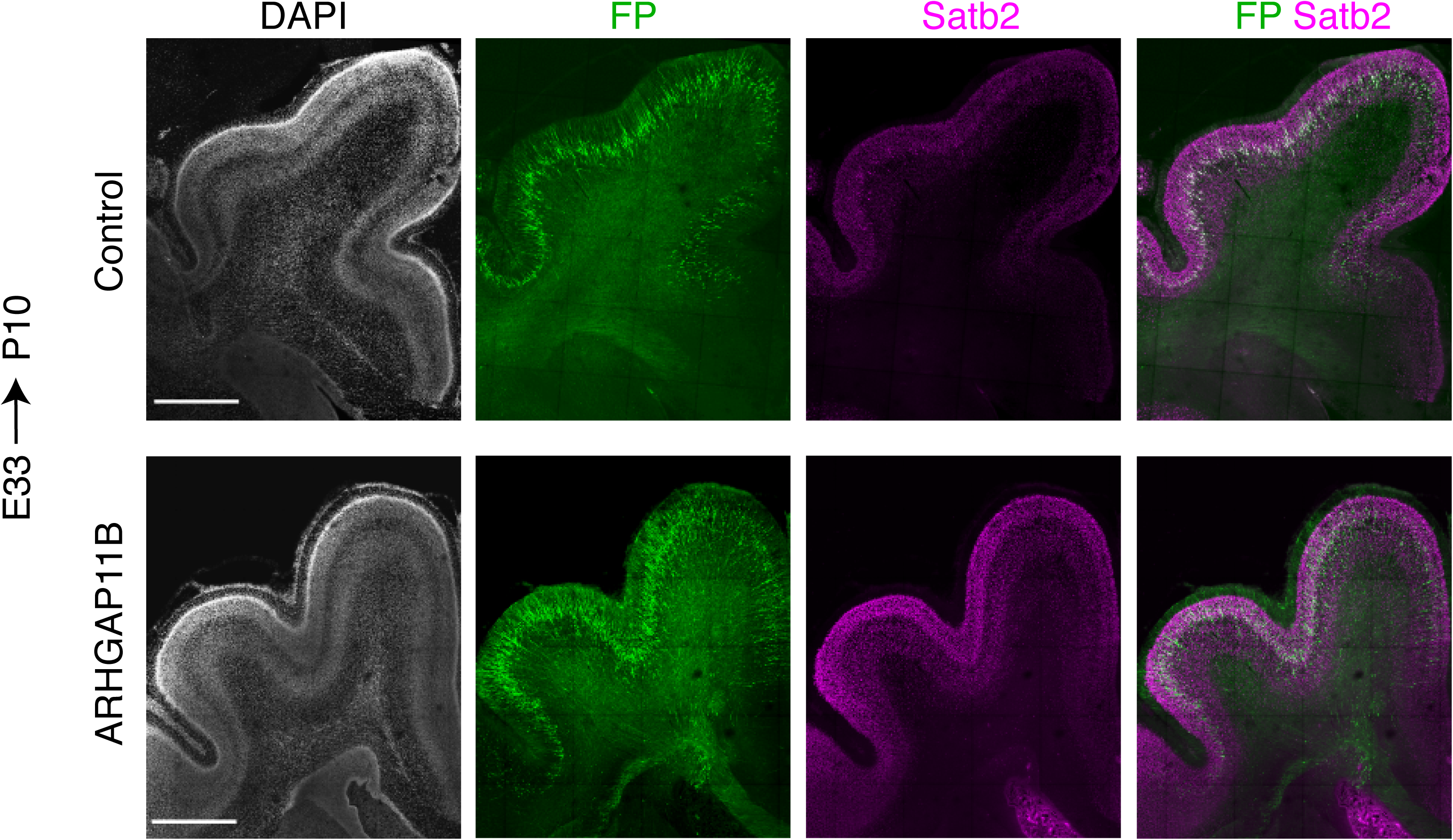
FP and Satb2 immunostaining patterns of control and ARHGAP11B-expressing developing ferret neocortex at P10. Ferret E33 neocortex was electroporated *in utero* with a plasmid encoding FP together with either a plasmid encoding ARHGAP11B or empty vector (Control), followed by double immunofluorescence for FP (green) and Satb2 (magenta), combined with DAPI staining (white), at P10. Images are maximum intensity projections of 5 optical sections. Scale bars, 1 mm.

**Figure 4 supplement 1.**
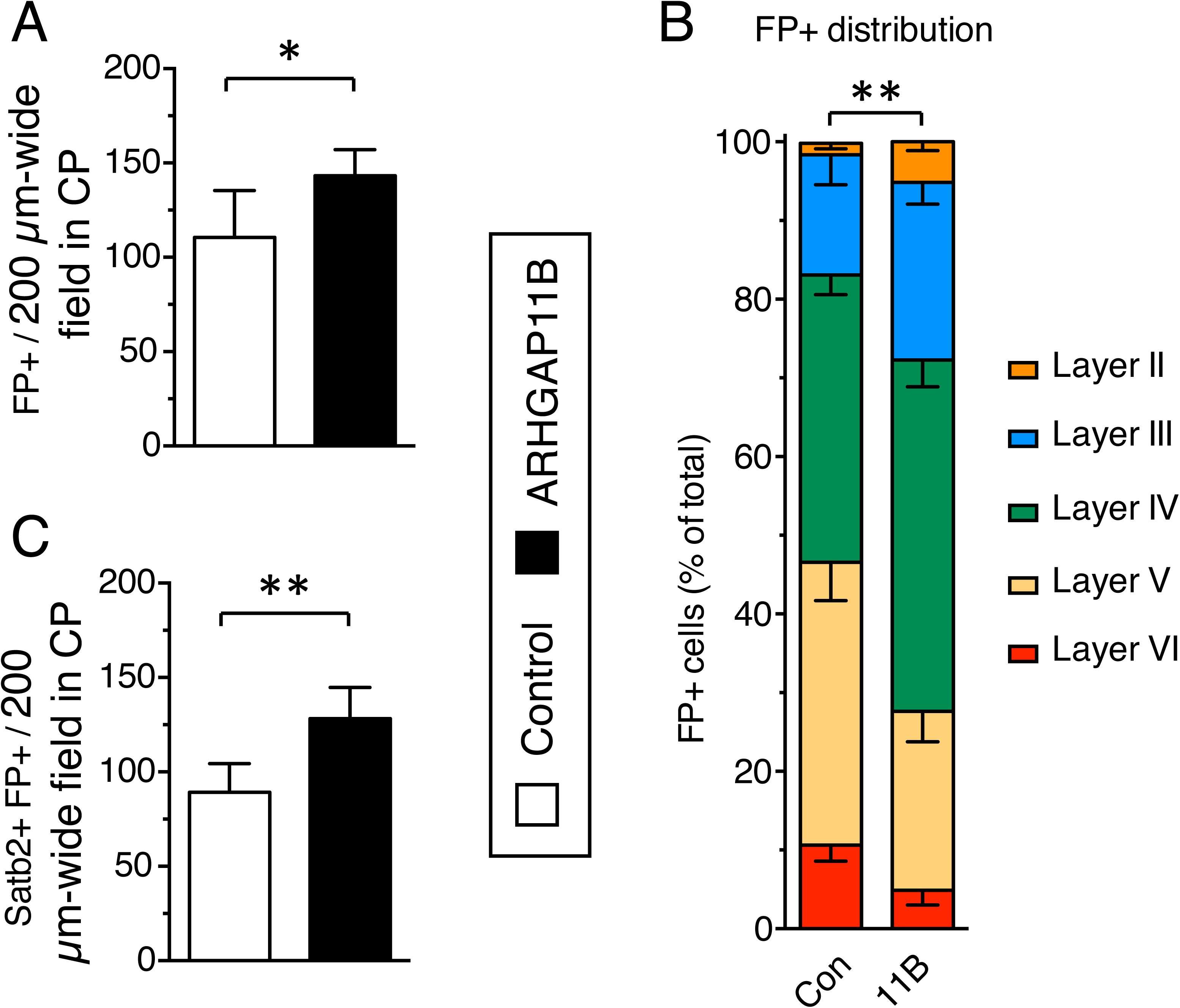
ARHGAP11B expression results in a greater abundance of FP+ cells and FP+ Satb2+ neurons in the CP. Ferret E33 neocortex was electroporated *in utero* with a plasmid encoding FP together with either a plasmid encoding ARHGAP11B or empty vector (Control), followed by analysis at P16. (A, C) Quantification of FP+ cells (A) and Satb2+ FP+ cells (C) in a 200 µm-wide field in the CP, upon control (white) and ARHGAP11B (black) electroporations. Data are the mean of 6 experiments. Error bars indicate SD; **, P <0.01; *, P <0.05, Student’s *t*-test. (B) Distribution of FP+ cells between the neuronal layers upon control (Con, left) and ARHGAP11B (11B, right) electroporations. Data are the mean of 6 experiments. Error bars indicate SD; **, P <0.01, two-way ANOVA with Bonferroni post-hoc tests (Layer V, Control vs. ARHGAP11B, P=0.014).

**Figure 4 supplement 2.**
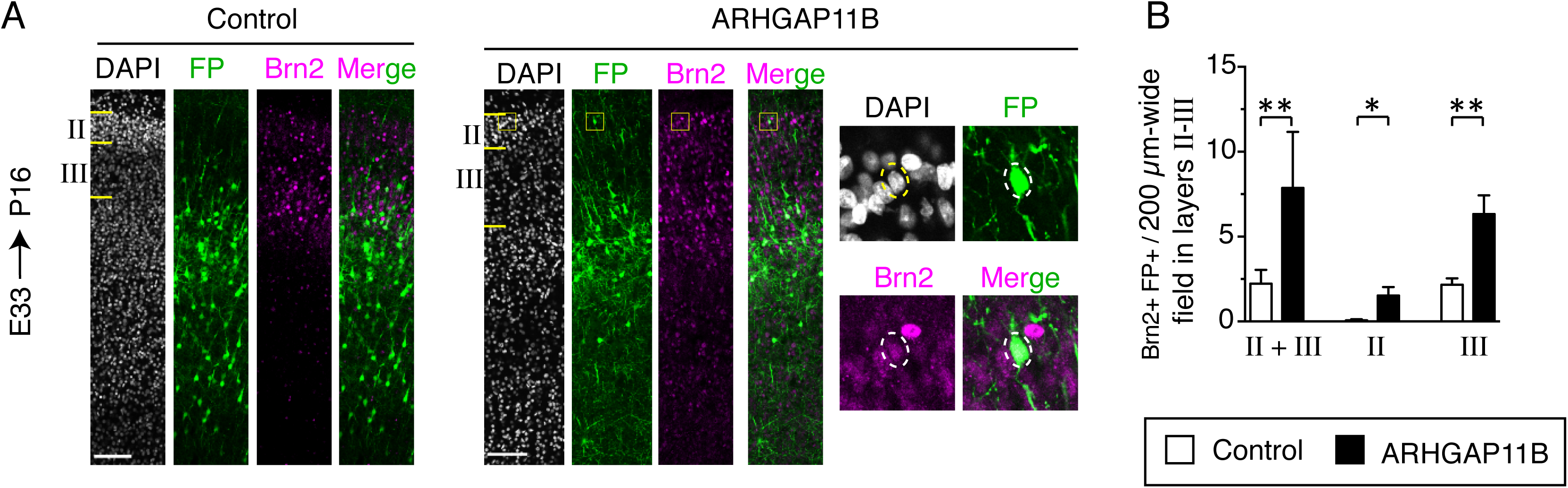
ARHGAP11B expression results in a greater abundance of FP+ Brn2+ neurons in cortical layers II and III. Ferret E33 neocortex was electroporated *in utero* with a plasmid encoding FP together with either a plasmid encoding ARHGAP11B or empty vector (Control), followed by double immunofluorescence for FP and Brn2, combined with DAPI staining, at P16. (A) Overview of the CP of the electroporated area showing the immunofluorescence for FP (green) and Brn2 (magenta) and the DAPI staining (white) (single optical sections). Layers II and III are indicated on the left. Scale bars, 100 μm. Boxes (50 × 50 μm) indicate a Brn2+ FP+ neuron in layer II, shown at higher magnification on the right. Dashed lines, cell body. (B) Quantification of Brn2+ FP+ cells in a 200 µm-wide field in layers II+III (left), in layer II only (center) and in layer III only (right), upon control (white) and ARHGAP11B (black) electroporations. Data are the mean of 5 experiments. Error bars indicate SD; **, P <0.01; *, P <0.05, Student’s *t*-test.

**Figure 4 supplement 3.**
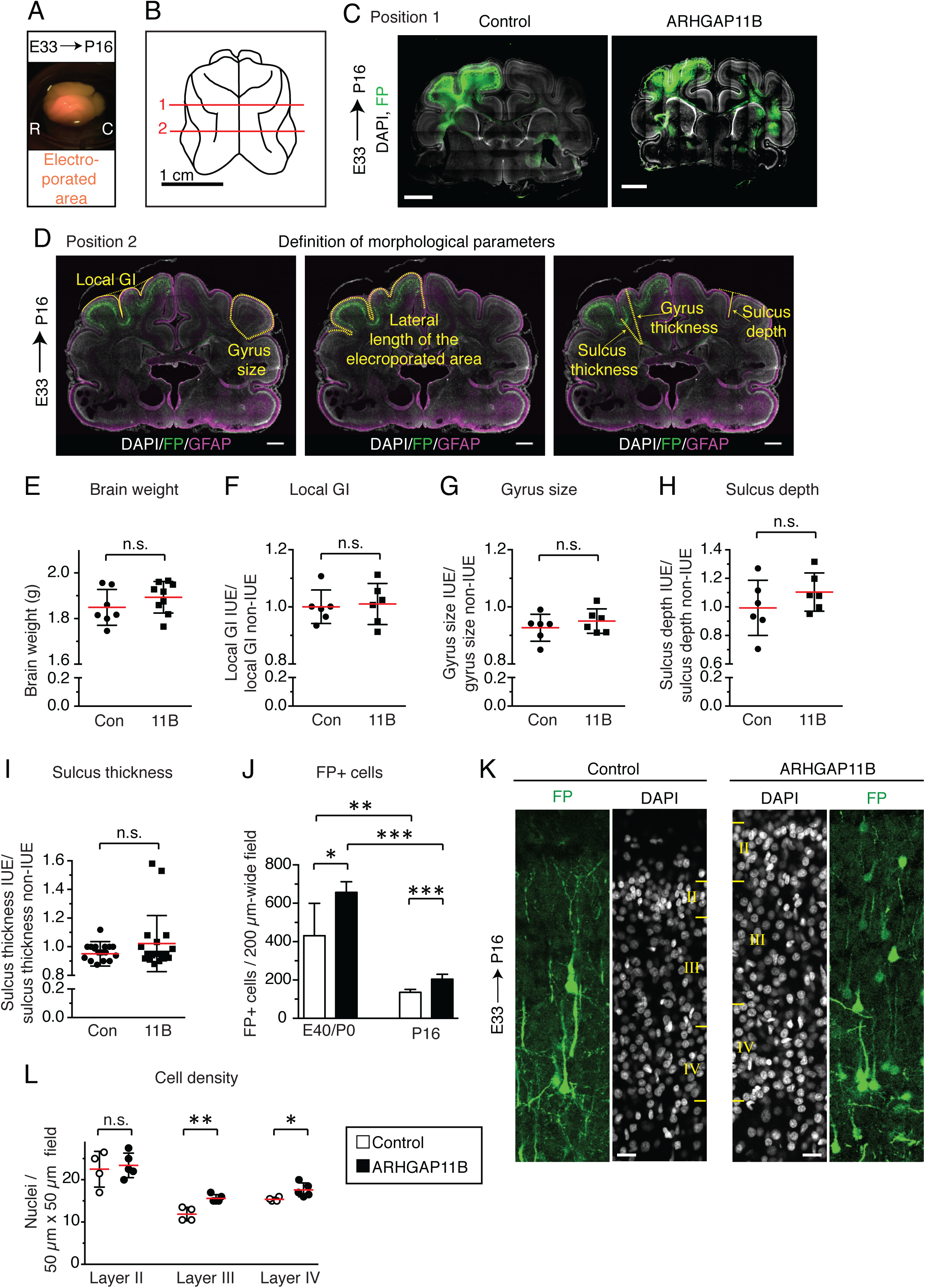
ARHGAP11B expression in developing ferret neocortex leads to its expansion. Ferret E33 neocortex was electroporated *in utero* with a plasmid encoding FP together with either a plasmid encoding ARHGAP11B or empty vector (Control), followed by analysis at E40/P0 (K, left) or P16 (all other panels). (A) Image of an electroporated brain, with the electroporated area in orange. R, rostral; C, caudal. (B) Schematic representation of the P16 ferret brain (dorsal view). Lines 1 and 2 indicate the positions at which all the morphological measurements were performed. Position 1 is exemplified in (C) and position 2 in (D). (C) Immunofluorescence for FP (green), combined with DAPI staining (white), of electroporated brains at position 1 (see B). Images are single optical sections. Scale bar, 1 mm. (D) Graphical definition of the measured morphological parameters. Double immunofluorescence for FP (green) and GFAP (magenta), combined with DAPI staining (white), of electroporated brain at position 2 (see B). Image (shown three times) is a single optical section. Yellow lines indicate measured parameters. See also Methods section for details. (E) Quantification of the weight of the control (Con) and ARHGAP11B-expressing (11B) ferret brains. Data are the mean (red lines) of 7 (control) or 9 (ARHGAP11B) brains. (F-I) Quantification of the indicated morphological parameters (see (D)), i.e. the gyrification index of the electroporated area (referred to as local GI) (F), gyrus size (G), sulcus depth (H) and sulcus thickness (I), of control (Con) and ARHGAP11B-expressing (11B) ferret neocortex. Measurements were performed at positions 1 and 2 (see (D)); for gyrus morphology, the posterior sigmoid gyrus (position 1), lateral gyrus (position 2) and the coronal gyrus (positions 1 and 2) were analyzed, yielding up to 4 data points; for sulcus morphology, the cruciate sulcus (position 1), lateral sulcus (position 2) and the suprasylvian sulcus (positions 1 and 2) were analyzed, also yielding up to 4 data points. Please refer to Sawada and Watanabe (2012) for the ferret gyri and sulci nomenclature. All data are expressed as ratio between electroporated hemisphere (IUE) and non-electroporated contra-lateral hemisphere (non-IUE). (F-H) Data are the mean (red lines) of 6 neocortices per condition. (I)Data are the mean (red lines) of 18 (control) and 19 (ARHGAP11B) sulci from 6 neocortices per condition. (J) Quantification of the number of FP+ cells in a 200 µm-wide field of cortical wall at E40/P0 (left) and P16 (right), upon control (white) and ARHGAP11B (black) electroporations. Data are the mean of 6 experiments. (K) Immunofluorescence for FP (green), combined with DAPI staining (white), of the indicated upper layers of the CP at position 2. Images are single optical sections. Scale bars, 20 μm. (L) Quantification of cell density, as revealed by DAPI staining of nuclei, in 50 μm × 50 μm fields of layer II (left), layer III (center) and layer IV (right) of the CP, upon control (white) and ARHGAP11B (black) electroporations. Data are the mean (red lines) of 4 (control) and 5 (ARHGAP11B) experiments, with 3 fields per layer per experiment. (E-J, L) Error bars indicate SD; ***, P<0.001; **, P <0.01; *, P <0.05; n.s., not statistically significant, Student’s *t*-test.

**Figure 4 supplement 4.**
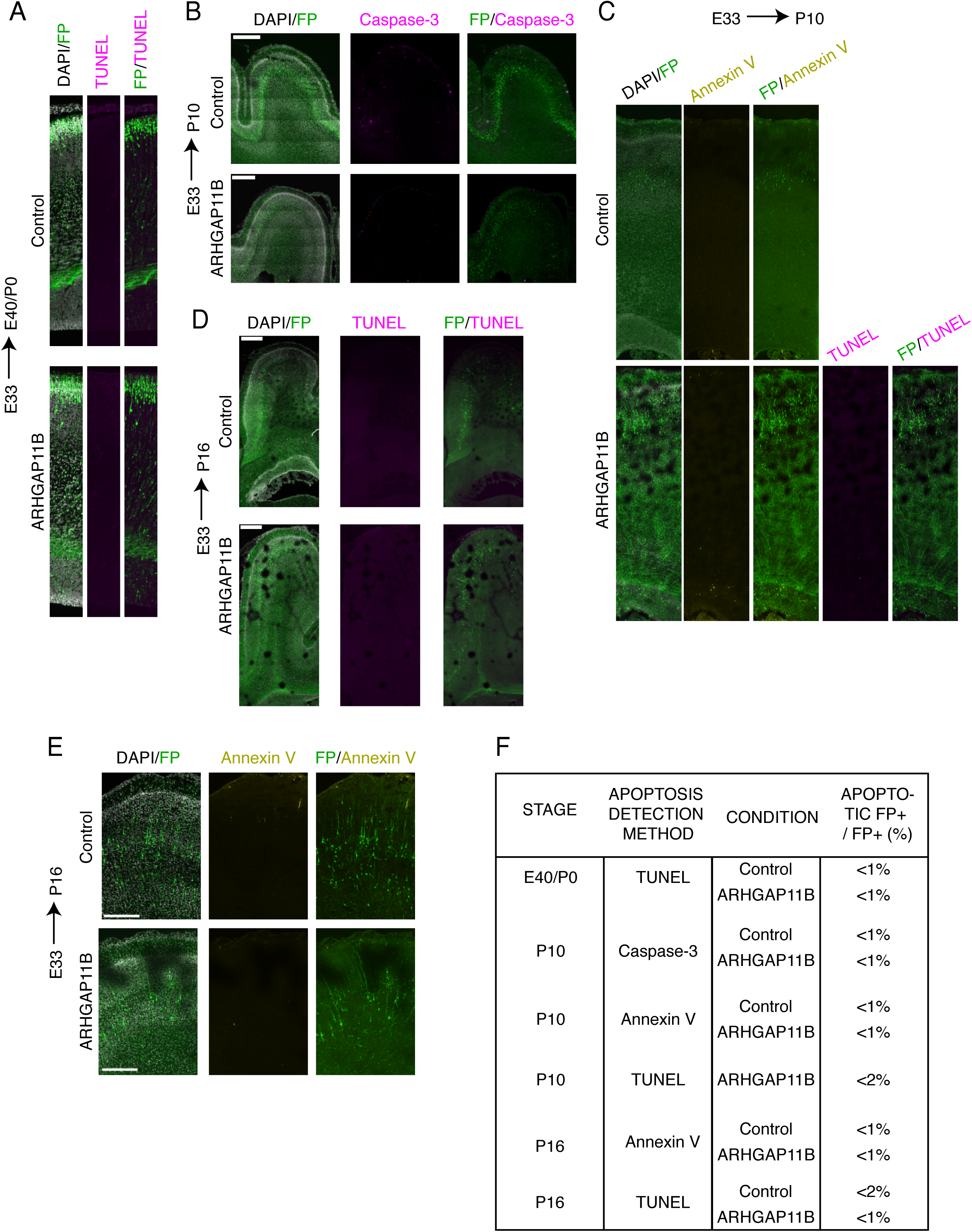
ARHGAP11B expression in developing ferret neocortex does not lead to increased cell death. Ferret E33 neocortex was electroporated *in utero* with a plasmid encoding FP together with either a plasmid encoding ARHGAP11B or empty vector (Control), followed by analysis at E40/P0 (A, F), P10 (B, C, F) and P16 (D, E, F). (A) Immunofluorescence for FP (green), combined with DAPI staining (white) and TUNEL (terminal deoxynucleotidyl transferase dUTP nick end labeling) staining (magenta), of the E40/P0 ferret neocortex. Image width, 200 μm. (B) Double immunofluorescence for FP (green) and Caspase-3 (magenta), combined with DAPI staining (white), of the P10 ferret neocortex. Scale bar, 500 μm. (C) Immunofluorescence for FP (green), combined with staining using fluorescently labeled annexin V (yellow) and DAPI staining (white), of the P10 ferret neocortex. TUNEL staining (magenta) is shown for the ARHGAP11B-expressing neocortex. Image width, 200 μm. (D) Immunofluorescence for FP (green), combined with DAPI staining (white) and TUNEL staining (magenta), of the P16 ferret neocortex. Scale bar, 500 μm. (E) Immunofluorescence for FP (green), combined with staining using fluorescently labeled annexin V (yellow) and DAPI staining (white), of the cortical plate of the P16 ferret neocortex. Scale bar, 300 μm. (A-E) All images are single optical sections. (F) Percentage of FP+ cells that are apoptotic. Note that at all stages analyzed and by all detection methods, ARHGAP11B-expressing neocortex does not show an increase in apoptosis. Data are the mean of at least two neocortices per condition.

**Figure 4 supplement 5.**
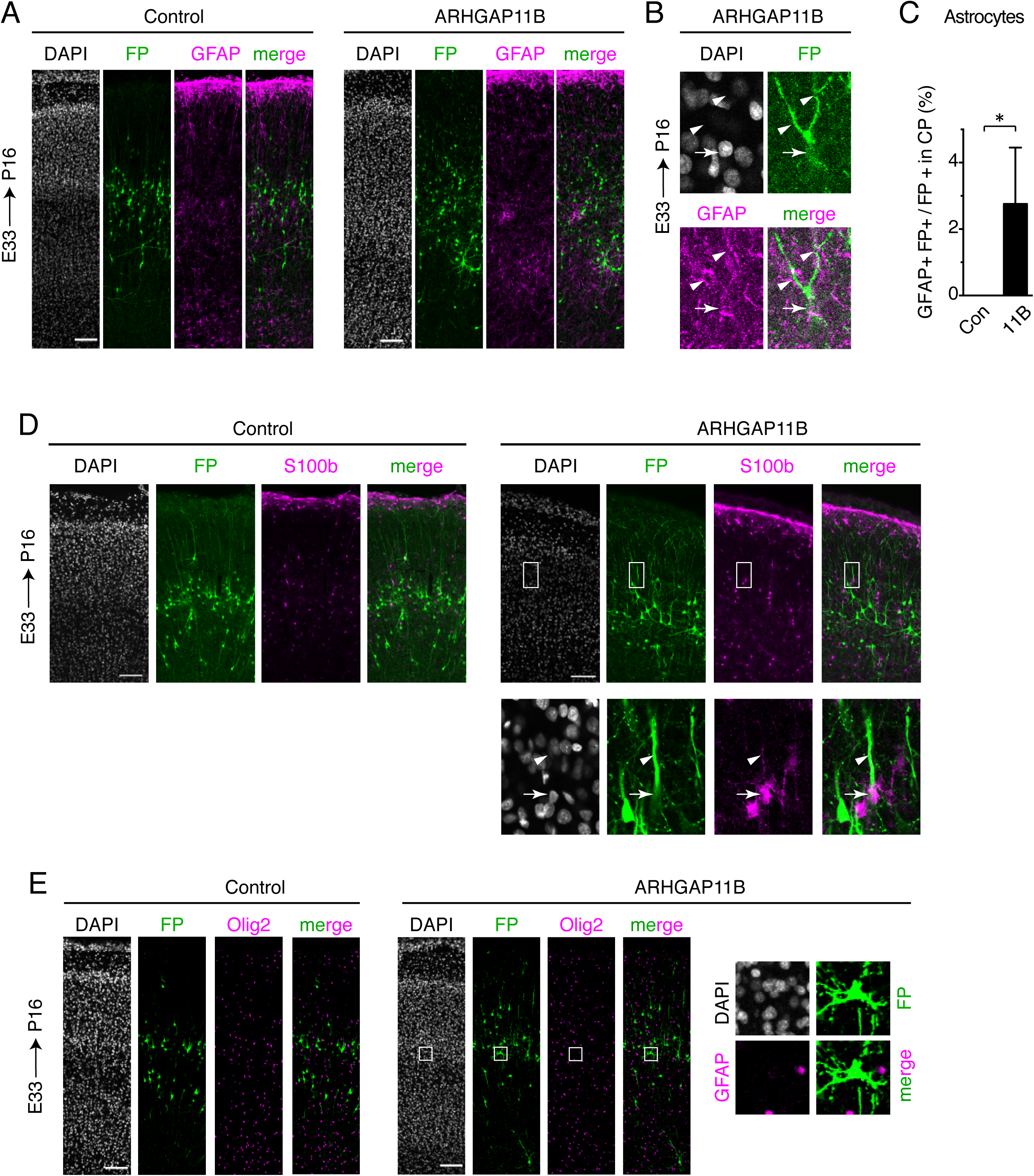
ARHGAP11B induces the appearance of astrocytes from the targeted progenitors in the developing ferret neocortex. Ferret E33 neocortex was electroporated *in utero* with a plasmid encoding FP together with either a plasmid encoding ARHGAP11B or empty vector (Control), followed by analysis at P16. (A, B) Double immunofluorescence for FP (green) and GFAP (magenta), combined with DAPI staining (white), of the CP (A, single optical sections). Scale bars, 100 μm. (B) High magnification of an FP+ GFAP+ ARHGAP11B-expressing cell in the CP from a different experiment than (A). Arrows, cell body; arrowheads, cell processes. Image width, 36.7 μm. (C) Percentage of FP+ cells in the CP that are GFAP+, upon control (white) and ARHGAP11B (black) electroporations. Data are the mean of 4 experiments. Error bars indicate SD; *, P<0.05, Student’s *t*-test. (D) Double immunofluorescence for FP (green) and S100ß (magenta), combined with DAPI staining (white), of the CP (single optical sections). Scale bars, 100 μm. Boxes (75.5 μm wide) indicate an FP+ S100ß+ cell in the CP, shown at higher magnification in the bottom images. Arrows, cell body; arrowheads, cell process. (E) Double immunofluorescence for FP (green) and Olig2 (magenta), combined with DAPI staining (white), of the CP (single optical sections). Scale bars, 100 μm. Boxes (52 × 52 μm) indicate an FP+ Olig2– cell in the CP, shown at higher magnification on the right. Note that no cells were found to be FP+ Olig2+.

**Supplemental Table 1.**
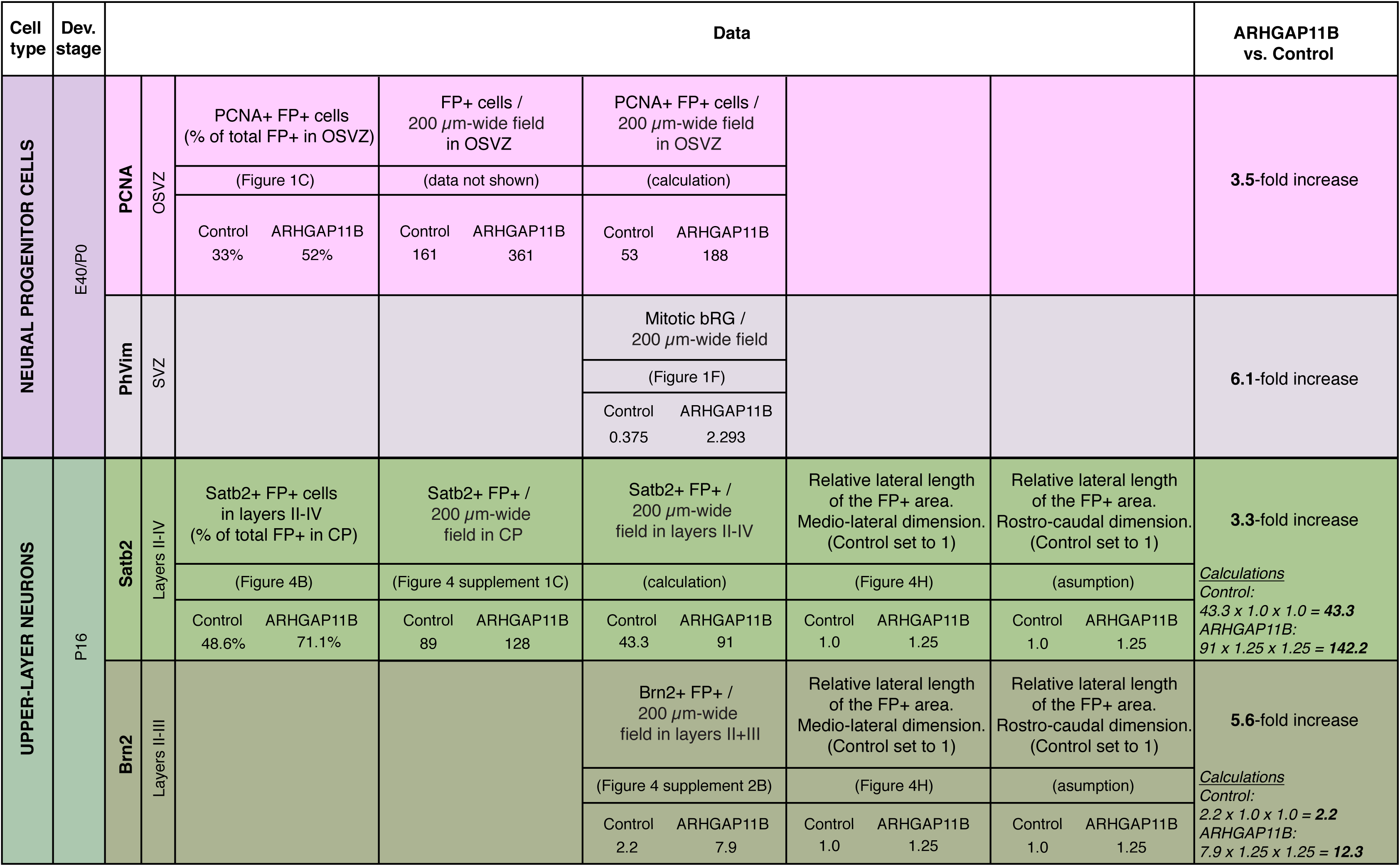
Overview of the effects of ARHGAP11B expression on neural progenitor cells and upper-layer neurons in developing ferret neocortex. Data taken from the indicated figure panels were used for the calculations shown. For the calculations pertaining to upper-layer neurons, it is assumed that the ARHGAP11B-induced increase in the lateral length of the FP+ area in the rostro-caudal dimension is equal to that in the medio-lateral dimension (1.25-fold).

## MATERIALS AND METHODS

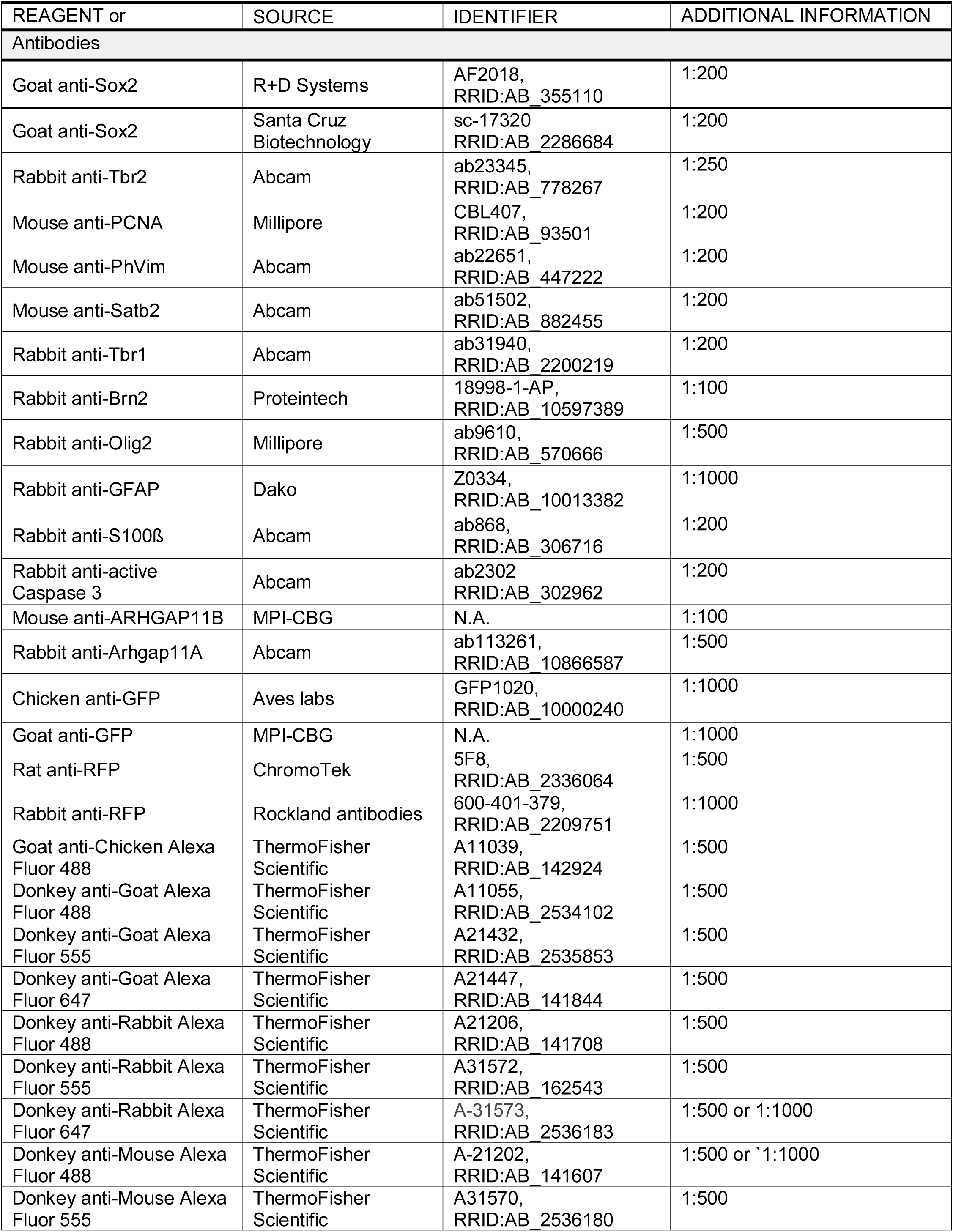
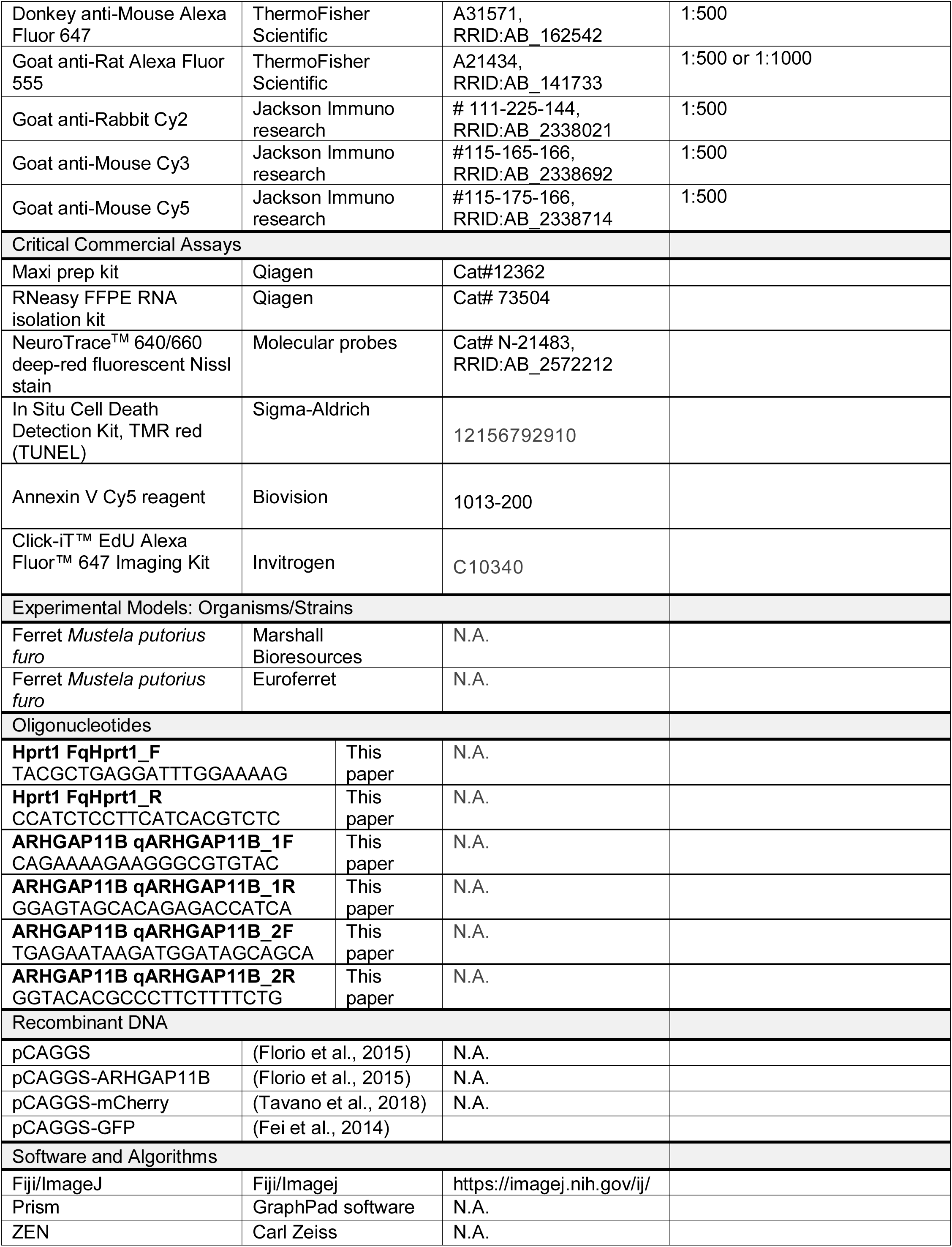
KEY RESOURCES TABLE

### Experimental animals

All experimental procedures were conducted in agreement with the German Animal Welfare Legislation after approval by the Landesdirektion Sachsen (licences TVV 2/2015 and TVV 21/2017). Timed-pregnant ferrets (*Mustela putorius furo*) were obtained from Marshall BioResources (NY, USA) or Euroferret (Copenhagen, Denmark) and housed at the Biomedical Services Facility (BMS) of MPI-CBG. Observed mating date was set to E0. Animals were kept in standardized hygienic conditions with free access to food and water and with an 16 h/8 h light/dark cycle. All experiments were performed in the dorsolateral telencephalon of ferret embryos, at a medial position along the rostro-caudal axis, in the prospective motor and somatosensory cortex.

### Plasmids

All plasmids used in this study were previously published (see the Key Resources table). All plasmids were extracted and purified using the EndoFree Plasmid Maxi kit (QIAGEN) following the manufacturer’s instructions.

### *In utero* electroporation of ferrets

*In utero* electroporation of ferrets was performed as originally established by Dr. Hiroshi Kawasaki with the modifications listed below (Kawasaki et al., 2012). Pregnant jills (with embryos at E33) were kept fasted for at least 3 h before the surgery. They were first placed in the narcosis box with 4% isoflurane. When in deep anesthesia, the ferrets were placed on the operation table and attached to the narcosis mask with a constant 3% isoflurane flow. Subsequently, the ferrets were injected subcutaneously with analgesic (0.1 ml Metamizol, 50 mg/kg), antibiotic (0.13 ml Synulox, 20 mg/kg or 0.1 ml amoxicilin, 10 mg/kg) and glucose (10 ml 5% glucose solution). A drop of Dexpanthenol Ointment solution was placed on their eyes to prevent eye dehydration during the surgical procedure. The ferret bellies were then shaved, sterilized with iodide and surgically opened. The uterus was exposed and embryos were injected intraventricularly with a solution containing 0.1% Fast Green (Sigma) in sterile PBS, 2.5 μg/μl of either pCAGGS vector (control) or pCAGGS-ARHGAP11B vector, in either case together with 1 μg/μl of pCAGGS vector encoding a fluorescent protein (pCAGGS-GFP or pCAGGS-mCherry). For embryonic studies where the position of the electroporated embryos in the uterus was known, the same fluorescent protein-encoding vector was co-electroporated together with either control or ARHGAP11B-expressing vectors. For postnatal studies, pCAGGS and pCAGGS-ARHGAP11B were co-electroporated with different fluorescent protein-encoding vectors to enable distinction of kits. For different experiments, these vectors were alternated. Electroporations were performed with six 50-msec pulses of 100 V at 1 sec intervals. Following electroporation, uterus was placed back in the peritoneal cavity and the muscle layer with the peritoneum were sutured using a 4-0 suture. The skin was sutured intracutaneously using the same thickness of the suture. Animals were carefully monitored until they woke up and then underwent postoperative care for the following 3 days (2 × daily 10 mg/kg amoxicilin, 3 × daily 25 mg/kg Metamizol).

### EdU labeling of ferret kits

For the neurogenic length experiments, a single pulse of EdU was injected intraperitoneally at P5, i.e. 13 days after *in utero* electroporations, and the animals were sacrificed 11 days later, at P16. The protocol for EdU injection was the same as the one previously published for ferret kits (Turrero Garcia et al., 2015).

### Isolation of ferret brains

Ferret embryos were isolated at E37 and E40. Ferret kits were sacrificed at P0, P10 and P16. For the isolation of embryos, pregnant jills underwent the second surgery that followed the same pre-operative care, anesthesia and analgesia as the first surgery. The sutures from the first operation were carefully removed and the uterus was exposed. A caesarian section was made and embryos were removed from the uterus. Subsequently a complete hysterectomy was performed, after which the muscle layer with peritoneum and skin were sutured and the animal underwent the same post-operative care as after the first surgery. Embryonic brains were isolated and fixed at 4°C for 48 h in 4% paraformaldehyde in 120mM phosphate buffer pH 7.4.

For the postnatal time points, the kits were sacrificed by intraperitonal injection of 4 mg/kg Xylazin + 40 mg/kg Ketamin. When in deep anaesthesia, the kits were perfused intracardially with PBS, followed by perfusion with 4% paraformaldehyde in 120mM phosphate buffer pH 7.4 at room temperature. Kit brains were then isolated and fixed for 48 h in 4% paraformaldehyde in 120 mM phosphate buffer pH 7.4 at 4°C.

Ferret jills whose kits were used postnatally also underwent the second surgery with the hysterectomy. Upon this surgery, they underwent the same post-operative care as after the first surgery. All jills were kept at the BMS of the MPI-CBG for at least two weeks after the second surgery. Afterwards they were donated for adoption. No adult ferrets were sacrificed in this study.

### Preparation of ferret brain slices

Upon fixation ferret brains were sectioned either on a vibratome or cryostat. Vibratome sections were 70 μm thick. Thickness of cryosections varied from 35 to 60 μm. Vibratome sections were either immunostained freshly prepared or conserved in cryoprotectant solution (30% sucrose, 30% ethylene glycol, 1% PVP40, 1.3 mM NaH_2_PO_4_, 3.9 mM Na_2_HPO_4_, 15 mM NaCl) and stored at –20°C for later use.

### Gene expression analysis by RT-qPCR

For analysis of *ARHGAP11B* expression, total RNA was isolated from two to three cryosections per developmental stage, using the RNeasy FFPE RNA isolation kit (Qiagen) including DNase-treatment following the manufacturer’s instructions. An additional DNase-treatment was performed on the isolated RNA using the DNA-free DNA Removal Kit (Life technologies). cDNA was synthesized using random hexamers and Superscript III Reverse Transcriptase (Life Technologies). qPCR was performed using the Light Cycler SYBR green Master mix (Roche) on a Light Cycler 96 (Roche). Gene expression data was normalized based on the housekeeping gene *Hprt1*. Primer sequences are provided in the Key Resources Table.

### Immunofluorescence

Immunofluorescence was performed as previously described (Kalebic et al., 2016). Antigen retrieval (1 h incubation with 10 mM citrate buffer pH 6.0 at 70°C in a water bath or oven) was performed for the following samples: all E37 samples, all E40/P0 samples, and vibratome sections of P10 and P16 samples which were used for immunostainings of the nuclear markers (Satb2, Brn2, Sox2, Olig2 and Ki67). When immunofluorescence for Tbr2 (E40/P0) was performed, a modified antigen retrieval protocol was used (1 h incubation with 10 mM citrate buffer pH 6.0 supplemented with 0.05% Tween-20 at 80°C in a water bath). Antigen retrieval was followed by three washes with PBS.

All the immunostainings, except the one for ARHGAP11B, were done as follows. Samples were subjected to permeabilization for 30 min in 0.3% Triton X-100 in PBS at room temperature, followed by quenching for 30 min in 0.1 M glycine in PBS at room temperature. Blocking was performed in a blocking solution (0.2% gelatin, 300 mM NaCl, 0.3% Triton X-100 in PBS) for 30 min. Primary antibodies were incubated in the blocking solution for 48 h at 4°C. Subsequently, the sections were washed three times in the blocking solution, incubated with secondary antibodies (1:500) and DAPI (Sigma) in the blocking solution for 1 h at room temperature, and washed again three times in the blocking solution before being either used for Nissl post-staining or directly mounted on microscopy slides with Mowiol.

For the immunostaining for ARHGAP11B, permeabilization was performed for 1 h in the 0.3% Triton X-100 solution at room temperature. Quenching was performed as for the other immunostainings. Blocking was done in 10% horse serum supplemented with 0.3% Triton X-100 (HS blocking solution) for 1 h. Primary antibodies were incubated in the same solution for 24 h, at 4°C. Subsequently, sections were washed five times in the HS blocking solution, incubated with the secondary antibodies (1:1000) and DAPI in the HS blocking solution for 1 h at room temperature, washed five times in PBS and mounted on microscopy slides with Mowiol. ARHGAP11B antibody used in this study is a mouse monoclonal antibody raised against a recombinant full length ARHGAP11B. As the first 220 amino acids of ARHGAP11B are 89% identical to the first 220 amino acids of the ferret Arhgap11a, we showed by immunostaining that the antibody we used did not recognize the endogenous ferret Arhgap11a (Figure 1 supplement 1E).

Staining with the Nissl fluorescent stain, EdU detection, annexin V labeling, and TUNEL labeling were performed after the antibody stainings. NeuroTrace™ 640/660 deep-red fluorescent Nissl stain (Molecular probes) was used, following the manufacturer’s instructions. EdU staining was performed using Click-iT™ EdU Alexa Fluor™ 647 Imaging kit (Invitrogen), following the manufacturer’s instructions. Annexin V labeling was performed using the Annexin V-Cy5 bright fluorescence reagent (BioVision), following the manufacturer’s instructions. TUNEL labeling was performed using the *In situ* cell death detection kit, ™R red (Sigma-Aldrich), following the manufacturer’s instructions.

### Image acquisition

Images of whole ferret brains (Figure 4 supplement 3A) were obtained as follows. Images were acquired using an iPhone 7 and taking a photography of the brain through a UV-protection filter mounted on a SZX 16 Olympus stereomicroscope, equipped with a fluorescence lamp.

Fluorescent images were acquired using a Zeiss LSM 880 upright single-photon point scanning confocal system. For the embryonic stages, images were taken as either 1-μm single optical sections with the 40x objective or 2-μm single optical sections with the 20x objective. For the postnatal stages, images were taken as either 2-μm single optical sections with the 20x objective or 7.2 μm single optical sections with the 10x objective. When the images were taken as tile scans, the stitching of the tiles was performed using the ZEN software. Subsequently, all images were analyzed and processed with ImageJ (http://imagej.nih.gov/ij/).

### Quantifications

All cell counts were performed in standardized microscopic fields using Fiji, processed using Excel (Microsoft), and results were plotted using Prism (GraphPad Software). For each condition, data (typically at least 3 microscopic fields) from one experiment (see definition below) were pooled, and the mean of the indicated number of experiments was calculated. Whenever possible the quantifications were done blindly.

The definition of the morphological parameters is depicted graphically in the Figure 4 supplement 3D. All the morphological parameters were quantified at two positions in the somatosensory cortex, as defined in Figure 4 supplement 3B, for the following gyri and sulci: posterior sigmoid gyrus, coronal gyrus, lateral gyrus, cruciate sulcus, suprasylvian sulcus and lateral sulcus (Sawada and Watanabe, 2012). All the morphological parameters are presented as a ratio of the value of the electroporated hemisphere and the value of the contralateral hemisphere, as established previously (Matsumoto et al., 2017). Calculations were as follows.

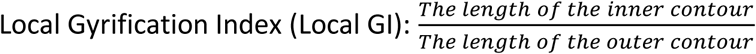

Local gyrification index is calculated as a ratio of the local GI of the electroporated area and the local GI of the equivalent area on the contralateral side. Gyrus size is calculated as a ratio of the size of an electroporated gyrus ad the size of the equivalent gyrus of the contralateral side. Sulcus depth is calculated as a ratio of the length of the line connecting the outer contour and the bottom of a sulcus and the equivalent line on the contralateral side. Sulcus thickness is calculated as a ratio of the thickness measured from the ventricular surface to the bottom of an electroporated sulcus and the equivalent thickness on the contralateral side. Gyrus thickness is calculated as a ratio of the thickness measured from the ventricular surface to the top of an electroporated gyrus and the equivalent thickness on the contralateral side. The upper layers thickness was measured as distance between the top of layer II and bottom of layer IV. Lateral length of the electroporated area was measured as the distance between the medial-most and the lateral-most cell on a coronal cross-section, following the inner contour of the neocortex. Cell density was measured as amount of DAPI+ nuclei in 50 µm × 50 μm fields.

### Statistical analysis

All statistics analyses were conducted using Prism (GraphPad Software). Sample sizes are reported in each figure legend, where the term “one experiment” would refer to one embryo for analysis at E37 or E40, and to one kit for postnatal analysis. Total number of litters analyzed was as follows: E37, 2 litters; E40/P0, 4 litters; P10, 2 litters; P16, 3 litters. Embryos or kits from all litters were included in the statistical analyses. Tests used were Two-way ANOVA with Bonferroni posttest, Student’s *t*-test and Mann-Whitney *U*-test. For each quantification, the statistical test and significance are indicated in the figure legend.

